# Gene-environment interactions in Multiple Sclerosis: a UK Biobank study

**DOI:** 10.1101/2020.03.01.971739

**Authors:** Benjamin Meir Jacobs, Alastair Noyce, Jonathan Bestwick, Daniel Belete, Gavin Giovannoni, Ruth Dobson

**Affiliations:** Preventive Neurology Unit, Wolfson Institute of Preventive Medicine, Barts and Queen Mary University of London; Royal London Hospital, Barts Health NHS Trust

## Abstract

**Importance:** Multiple Sclerosis (MS) is a neuro-inflammatory disorder caused by a combination of environmental exposures and genetic risk factors. We sought to determine whether genetic risk modifies the effect of environmental MS risk factors.

**Methods:** People with MS were identified within UK Biobank using ICD10-coded MS or self-report. Associations between environmental risk factors and MS risk were quantified with a case-control design using multivariable logistic regression. Polygenic risk scores (PRS) were derived using the clumping-and-thresholding approach with external weights from the largest genome-wide association study of MS. Separate scores were created including (PRS_MHC_) and excluding (PRS_Non-MHC_) the MHC locus. The best performing PRS were identified in 30% of the cohort and validated in the remaining 70%. Interaction between environmental and genetic risk factors was quantified using the Attributable Proportion due to interaction (AP) and multiplicative interaction.

**Results:** Data were available for 2250 people with MS and 486,000 controls. Childhood obesity, earlier age at menarche, and smoking were associated with MS. The optimal PRS were strongly associated with MS in the validation cohort (PRS_MHC_: Nagelkerke’s Pseudo-R^2^ 0.033, p=3.92×10^−111^; PRS_Non-MHC_: Nagelkerke’s Pseudo-R^2^ 0.013, p=3.73×10^−43^). There was strong evidence of interaction between polygenic risk for MS and childhood obesity (PRS_MHC_: AP=0.17, 95% CI 0.06 - 0.25, p=0.004; PRS_Non-MHC_: AP=0.17, 95% CI 0.06 - 0.27, p=0.006).

**Conclusions and Relevance:** This study provides novel evidence for an interaction between childhood obesity and a high burden of autosomal genetic risk. These findings may have significant implications for our understanding of MS biology and inform targeted prevention strategies.

## Introduction

Susceptibility to Multiple Sclerosis (MS) is multifactorial with genetic and environmental determinants. ^1,2,3^. Environmental exposures associated with MS risk include smoking, solvent exposure, childhood obesity, vitamin D deficiency, increasing latitude, and infectious mononucleosis (IM) ^2,3^. The largest genome-wide association study (GWAS) of MS risk performed by the International Multiple Sclerosis Genetic Consortium (IMSGC) revealed 233 independent signals which account for ∼48% of the estimated heritability of MS^1^. Attempts to model MS risk using polygenic risk scores have had some success^4–6^, supporting the view that MS susceptibility is influenced by common variants across the genome, in addition to the contribution from the Major Histocompatibility Complex (MHC).

A large proportion of MS risk remains unexplained despite the well-described genetic architecture^1^. One potential explanation for this ‘missing risk’ may is the presence of gene-environment interactions, whereby the effect of certain genes or variants may depend on exposure to environmental risk factors.

Evidence from Scandinavian and North American cohorts suggests that environmental influences on MS risk can be modified by HLA genotype. The deleterious effects of childhood obesity, smoking, IM, and solvent exposure on MS risk are potentiated among carriers of the HLA DRB1*15 allele and those lacking the protective HLA A*02 genotype^7–10^. It is not currently known whether gene-environment interactions in MS extend beyond the HLA locus^11,12^.

In this work, we harnessed the power of UK Biobank to extend our understanding of how common genetic variation interacts with environmental factors associated with MS development. We achieved this by firstly performing a large case-control study to confirm the role of established risk factors in this cohort, and by developing and validating polygenic risk scores (PRS) for MS which both included and excluded the MHC. Finally, we used these data to look for potential interactions between polygenic risk and environmental factors associated with MS.

## Methods

### Data sources

UK Biobank is a longitudinal cohort study described in detail elsewhere^13^. In brief, participants between the ages of 40 and 69 were recruited between 2006 and 2010 from across the UK. Participants underwent genotyping, donated body fluid samples, and answered a range of questions about lifestyle, environmental and demographic factors. Health records were linked to participants using Hospital Episode Statistics. Phenotype data are composed of survey data, linked healthcare records, anthropometric measurements, and a variety of other biochemical and imaging data (which was not used in this study).

### Identification of cases and controls

Cases were defined by ICD-coded diagnoses (ICD10-G35; ICD9-3409), self-reported MS diagnosis, or a GP-coded diagnosis. Age at diagnosis was determined using the first recorded MS diagnostic code (see supplementary methods for further details). Controls were unmatched UK Biobank participants without a coded diagnosis of MS. Individuals diagnosed with MS prior to age 20 were excluded due to concerns around the timing of exposures. Participant flow through the study is depicted in supplementary figure 1; diagnostic codes used are provided in supplementary data.

### Genotype data

Genotyping and quality control protocols are described in detail elsewhere^14^. Imputed HLA alleles were provided by UK Biobank. HLA alleles were imputed to four-digit resolution using the HLA*IMP:02 software with a multi-population reference panel (see https://biobank.ctsu.ox.ac.uk/crystal/crystal/docs/HLA_imputation.pdf). We extracted each participant’s allelic dosage for the MS risk allele HLA-DRB1*15:01 and the protective allele HLA-A*02:01 by thresholding posterior allele probabilities at 0.7 as suggested by UK Biobank. These two HLA alleles were used as they have the largest effect sizes across multiple studies^2^. Genetic principal components were supplied by UK Biobank (field ID 22009).

### Definition of exposures

Exposures were selected if they pertained to early life/adolescence (to mitigate the risk of reverse causation), and were previously associated with MS in at least one other observational cohort. Selected exposures were captured from baseline data recorded in UKB, along with age, ethnicity, sex, birth latitude, and Townsend deprivation index at recruitment (supplementary table 1).

We examined the following ten early life/environmental exposures: month of birth, having been breastfed as a child, childhood body size at age 10 (CBS_10_, a proxy for childhood obesity), exposure to maternal smoking, age at menarche (females), age at voice breaking (males), age at first sexual intercourse, smoking status prior to age 20, birth weight, and infectious mononucleosis prior to age 20. Where multiple data points were available for a participant, the first recorded reading was used.

Childhood body size was dichotomised and participants were classified as “not overweight” if they answered “thinner” or “average”, and “overweight” if they answered “plumper”. Smoking status was characterised as “ever” or “never” smoking. Age at menarche was treated as a continuous variable and analyses regarding menarche were restricted to women. IM status prior to age 20 was defined using the source of first report fields. Participants whose IM diagnosis was reported after age 20 were coded as having not had IM. Vitamin D status was not included, as vitamin D levels are only available from the initial visit (i.e. at study recruitment), which in the majority of cases was subsequent to diagnosis.

### Case-control study

For each risk factor, we built a multivariable logistic regression model modelling MS status as the outcome, with age, sex, ethnicity, current deprivation status, and birth latitude as potential confounding covariates^15^.

The strength of evidence for association with MS was determining using the model likelihood ratio, comparing the full model to a null model comprising only the confounding covariates. Strong evidence for association was defined using a Bonferroni-adjusted p-value threshold to maintain an alpha of 0.05 (p_threshold_=0.05/10=0.005). Risk factors robustly associated with MS at alpha<0.05 were then combined in a multivariable model including the most potent genetic risk factors, HLA DRB1*15:01 and HLA A*02:01, to assess whether their effects showed evidence of independent association with MS.

### Development of polygenic risk scores (PRS) for MS

A variety of PRS were created using the clumping-and-thresholding approach with external weights derived from the IMSGC discovery stage meta-analysis (supplementary methods). We created scores both including the MHC region (PRS_MHC_) and excluding this region (PRS_Non-MHC_). To validate the PRS, the dataset was divided randomly into a training set (30%, n_MS_=589, n_control_=112,724) and a testing set (70%, n_MS_=1237, n_control_=263,159, supplementary figure 1). To determine the optimal PRS, we constructed multivariable logistic regression models for each PRS with MS status as the outcome with age, sex, Townsend deprivation index, and the first four genetic principal components (PCs) as confounding covariates.

PRS performance was evaluated using Nagelkerke’s Pseudo-R^2^ metric, which is analogous to the R^2^ derived from linear regression models. Nagelkerke’s Pseudo-R^2^ was calculated comparing the full model including the PRS to a null model comprising the confounding covariates only. This procedure was repeated for all 64 scores (supplementary table 4). Altering the number of PCs adjusted for did not substantially alter the results (supplementary figures 5&6). Further validation is described in the supplementary methods.

### PRS x Environment interactions

The optimal PRS_MHC_ and PRS_Non-MHC_ were used to look for evidence of genome-wide gene-environment interactions using exposures identified as significantly associated with MS in the case-control study. All interaction analyses were conducted in the testing set to avoid PRS overfitting. Interaction was assessed on the additive and multiplicative scales (supplementary methods for full details). Multiplicative interaction was quantified using the interaction term beta from logistic models, and additive interaction was quantified using the Attributable Proportion due to interaction (AP).

### HLA x PRS interactions

To determine whether non-MHC genetic risk of MS modulates the effects of the most potent MHC risk allele, DRB1*15:01, we calculated additive and multiplicative interaction statistics using the methods described above, considering both the DRB 1*15:01 genotype (dominant-coding) and the non-MHC PRS as independent covariates.

### Association of PRS with disease measures

To determine whether the MS-PRS was associated with age at first report and disability status, we constructed regression models in the testing set. For age at first MS diagnostic code report, values were normalised using the inverse-rank normalisation. Linear regression models were constructed, using age, sex, Townsend score, and the first four PCs as covariates. Disability status was assessed using the UKB field ‘Attendance/disability/mobility allowance’ (field 6146), and recoded this as a binary variable (i.e. participants were coded as ‘1’ if they claimed any of the blue badge, attendance allowance, or disability living allowance, and as ‘0’ if not). Logistic regression models were then constructed using the same covariates as above (age, sex, Townsend score, first four genetic PCs).

### Ethical approval

This work was performed using data from UK Biobank (REC approval 11/NW/0382). All participants gave informed consent on Biobank registration and are free to withdraw from the study at any point, at which point their data are censored and cannot be included in further analyses.

### Computing

This research was supported by the High-Performance Cluster computing network hosted by Queen Mary, University of London^19^. Statistical analyses were performed in R version 3.6.1. Extraction of European individuals from the 1000 genomes reference genome was conducted using vcftools. Construction of the polygenic risk score, application of the polygenic risk score to individuals, and quality control were performed in PLINK 1.9 and PLINK2. All code used in this study is available on GitHub (@benjacobs123456).

## Results

### Population demographics

Phenotype and genotype data were available for 488,276 UK Biobank participants comprising 2276 people with MS and 486,000 unmatched controls. The median age at first MS report was 43.5 (IQR 16.1, supplementary figure 2). Demographic characteristics are shown in table 1. Characteristics of individuals with MS were consistent with published observational data (72.7% female, 98.1% White British).

**Table 1:**
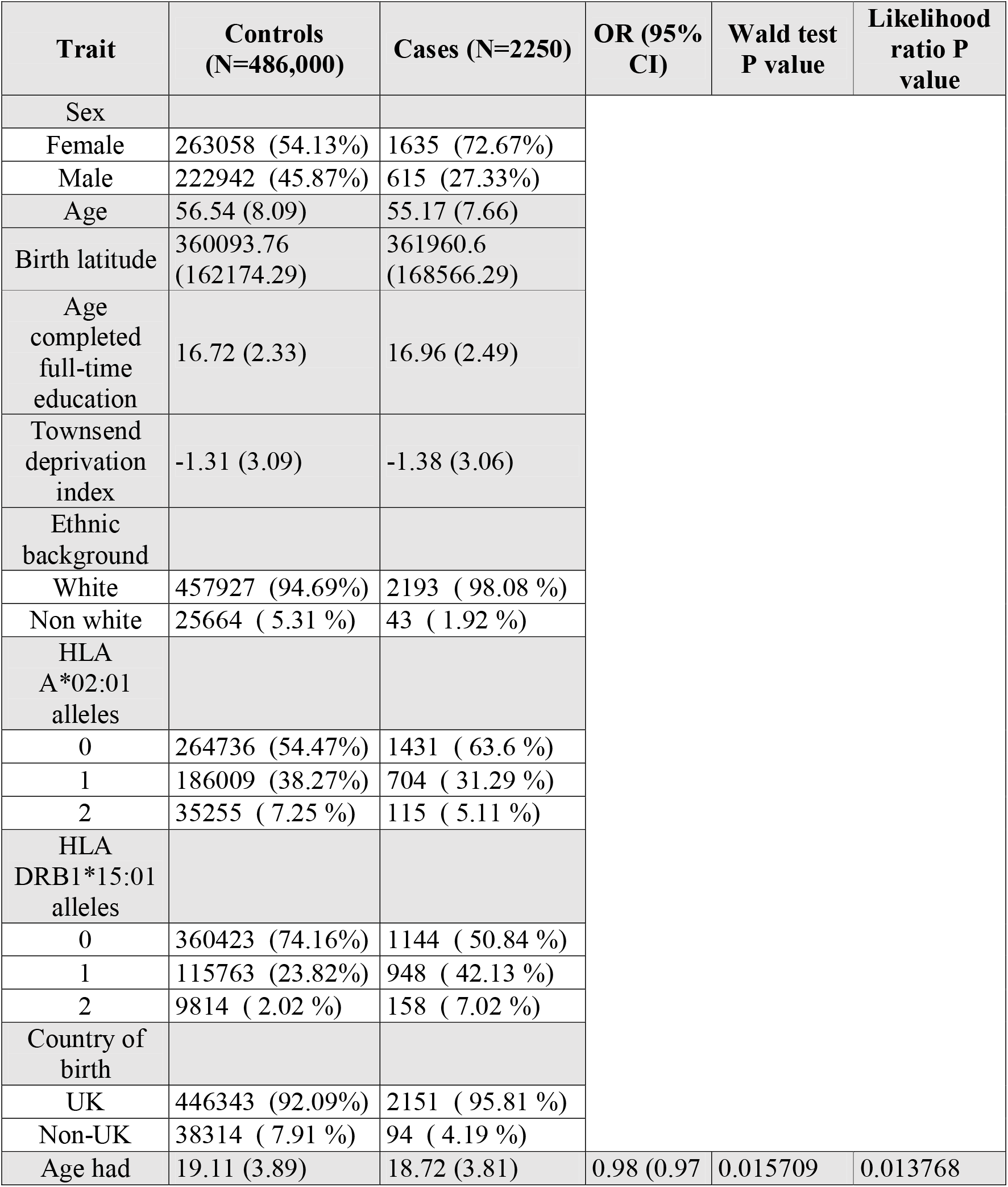
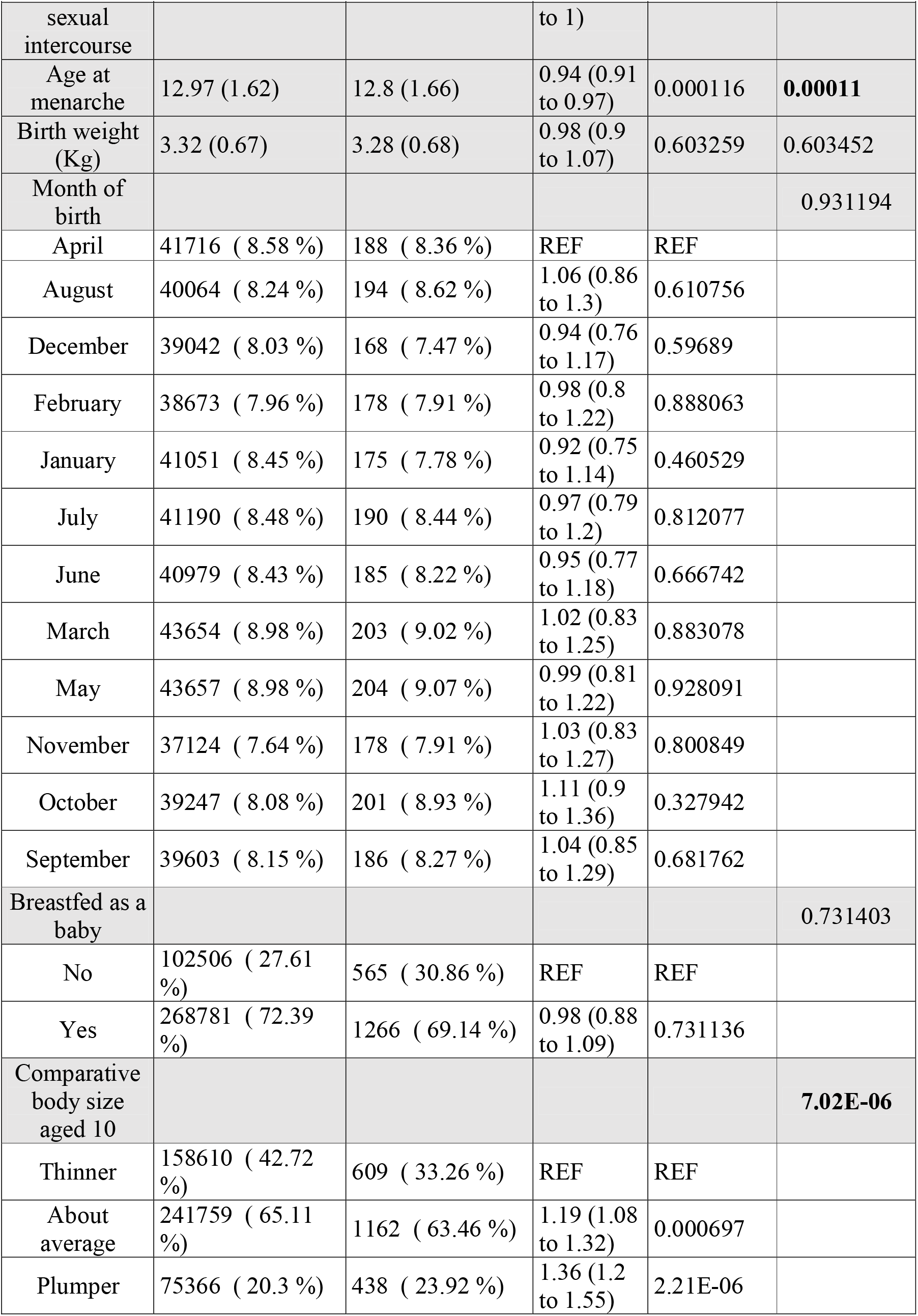
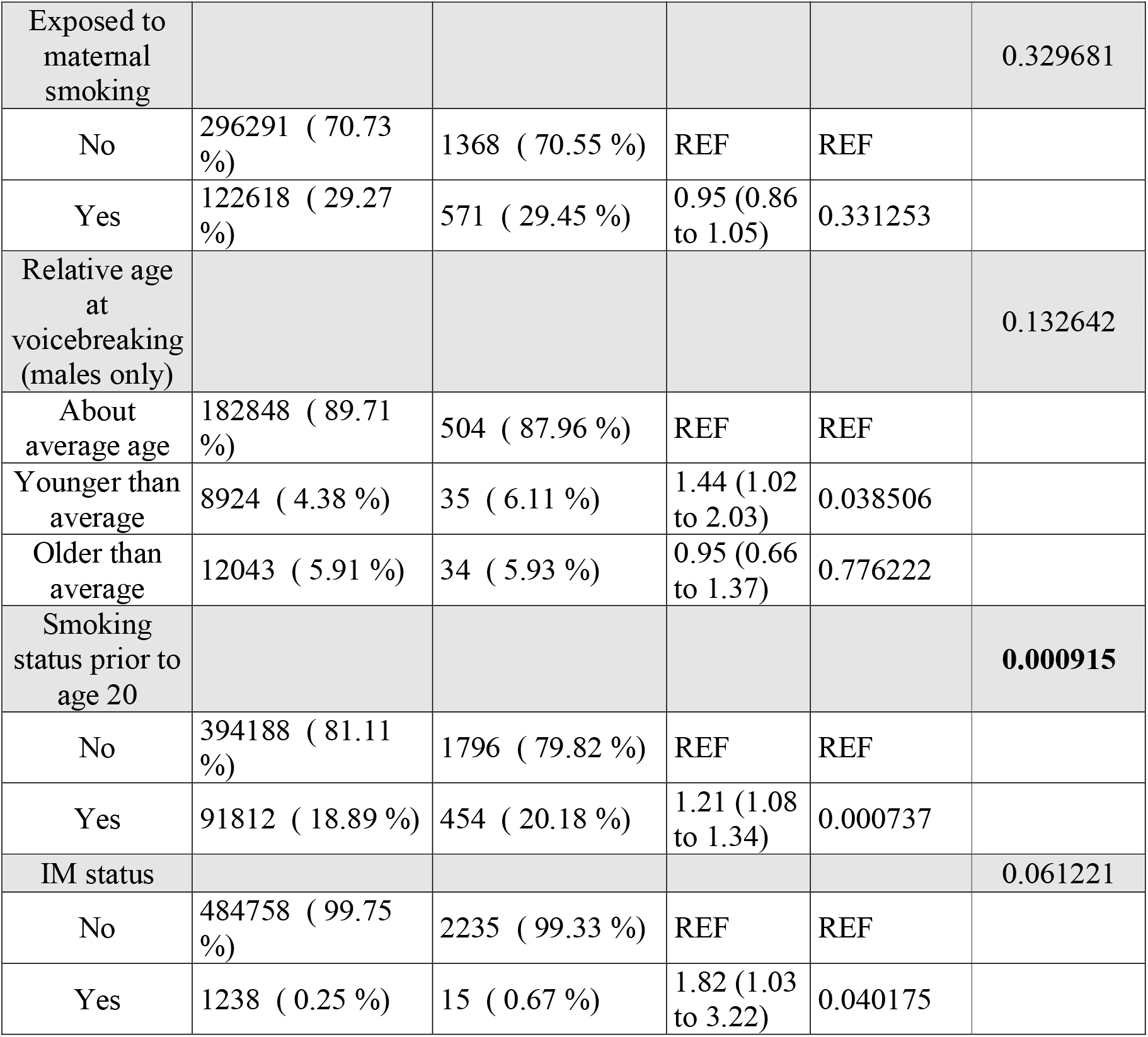
demographic characteristics of included participants and results from the case-control study. Continuous variables are presented as mean(SD), categorical variables are presented as n(%). Missing data are not tabulated. Proportions are calculated as a proportion of individuals with non-missing data for each variable. Column 4 shows odds ratios (ORs) and 95% confidence intervals (CIs) for Multiple Sclerosis for each exposure studied. Odds ratios represent the output of multivariable logistic regression models incorporating age, sex, ethnicity, birth latitude and current deprivation as covariates. Wald test P values represent the test of the null hypothesis beta = 0 for each term, whereas likelihood ratio P values represent the overall model fit compared to a null model comprising only confounding covariates. P values exceeding the Bonferroni multiple testing threshold (alpha=0.05) are shown in bold. Reference values for categorical covariates are denoted as ‘REF’.

### Exposures associated with MS in UK Biobank

There was strong evidence for association between three of the ten risk factors examined and MS (P_Bonf_<0.05): higher childhood body size at age 10 (‘plumper than average’ vs ‘thinner than average’: OR 1.36, 95% CI 1.20 - 1.55), smoking prior to age 20 (OR 1.21, 95% CI 1.08 - 1.34), and earlier menarche (OR 0.94, 95% CI 0.91 - 0.97, figure 1, table 1). The effects of these three risk factors remained similar in a combined model incorporating HLA DRB1*15:01 and HLA A*02:01 genotype (supplementary table 3).

**Figure 1:**
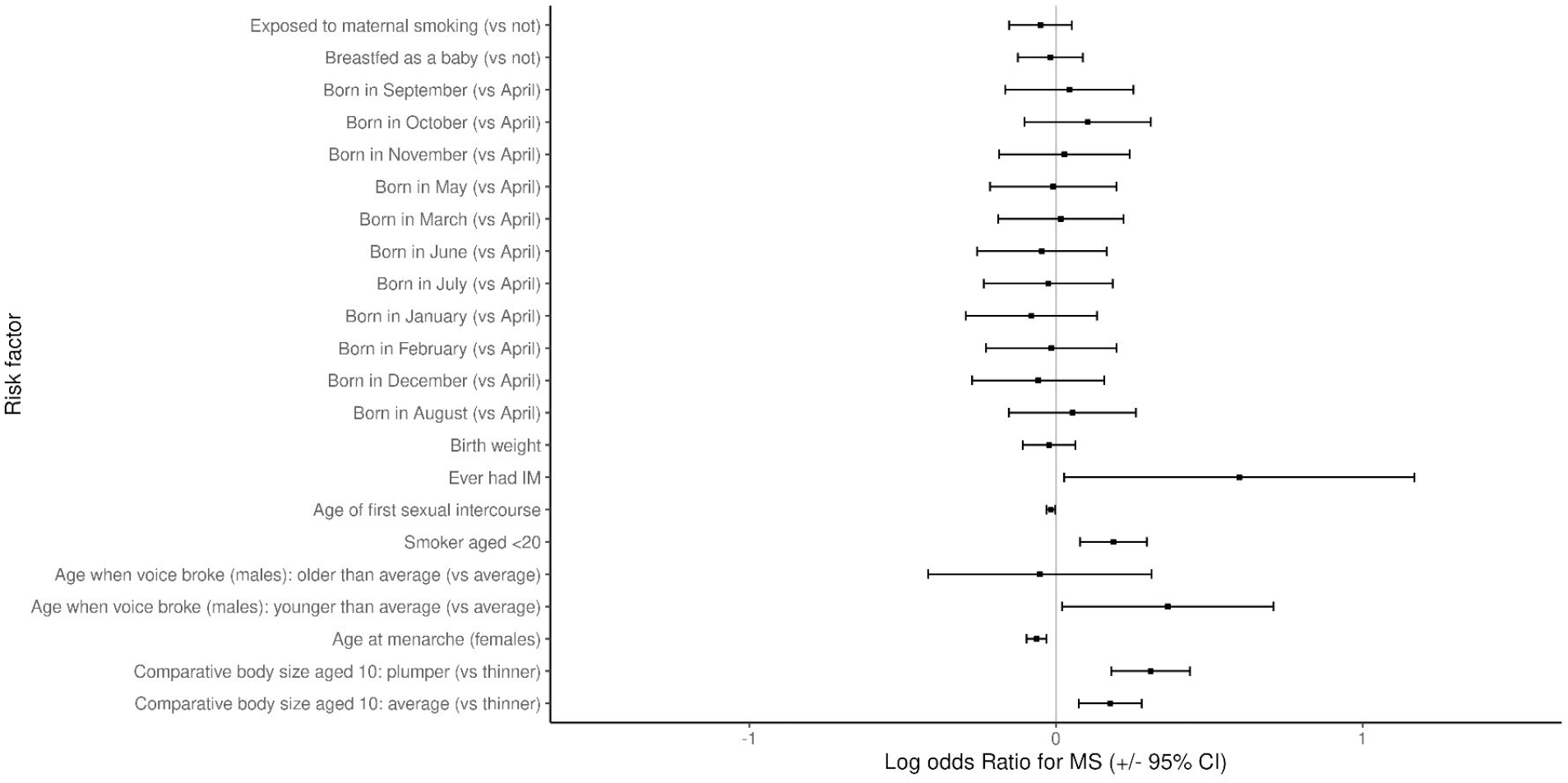
odds ratios and 95% confidence intervals for the association of each exposure with MS. ORs and CIs are from the output of a multivariable logistic regression with the following covariates: age, sex, ethnicity, birth latitude, current deprivation status, and the exposure in question. For menarche (females only) and voice-breaking (males-only), sex was not included as a covariate.

### Development and validation of PRS for MS

The optimal PRS_MHC_ and PRS_Non-MHC_ explained 3.5% and 1.3% of MS risk in the training set respectively (figure 2a, supplementary table 4, supplementary figures 5 and 6). Both scores were strongly associated with MS in the testing set (PRS_MHC_: Nagelkerke’s Pseudo-R^2^ 0.033, p=3.92×10^−111^; PRS_Non-MHC_: Nagelkerke’s Pseudo-R^2^ 0.013, p=3.73×10^−43^, figure 2, supplementary table 5). Both scores were reasonably well-calibrated (figure 3a) with good discriminative performance (AUC_MHC_ 0.71, AUC_non-MHC_ 0.67, AUC_null_ 0.63; figure 3b). There was no evidence of association between the PRS_MHC_ and PRS_Non-MHC_ and age at MS report (figure 3c & 3d) or claiming of disability benefits (p_MHC_=0.44, p _Non-MHC_=0.96, supplementary figure 7).

**Figure 2:**
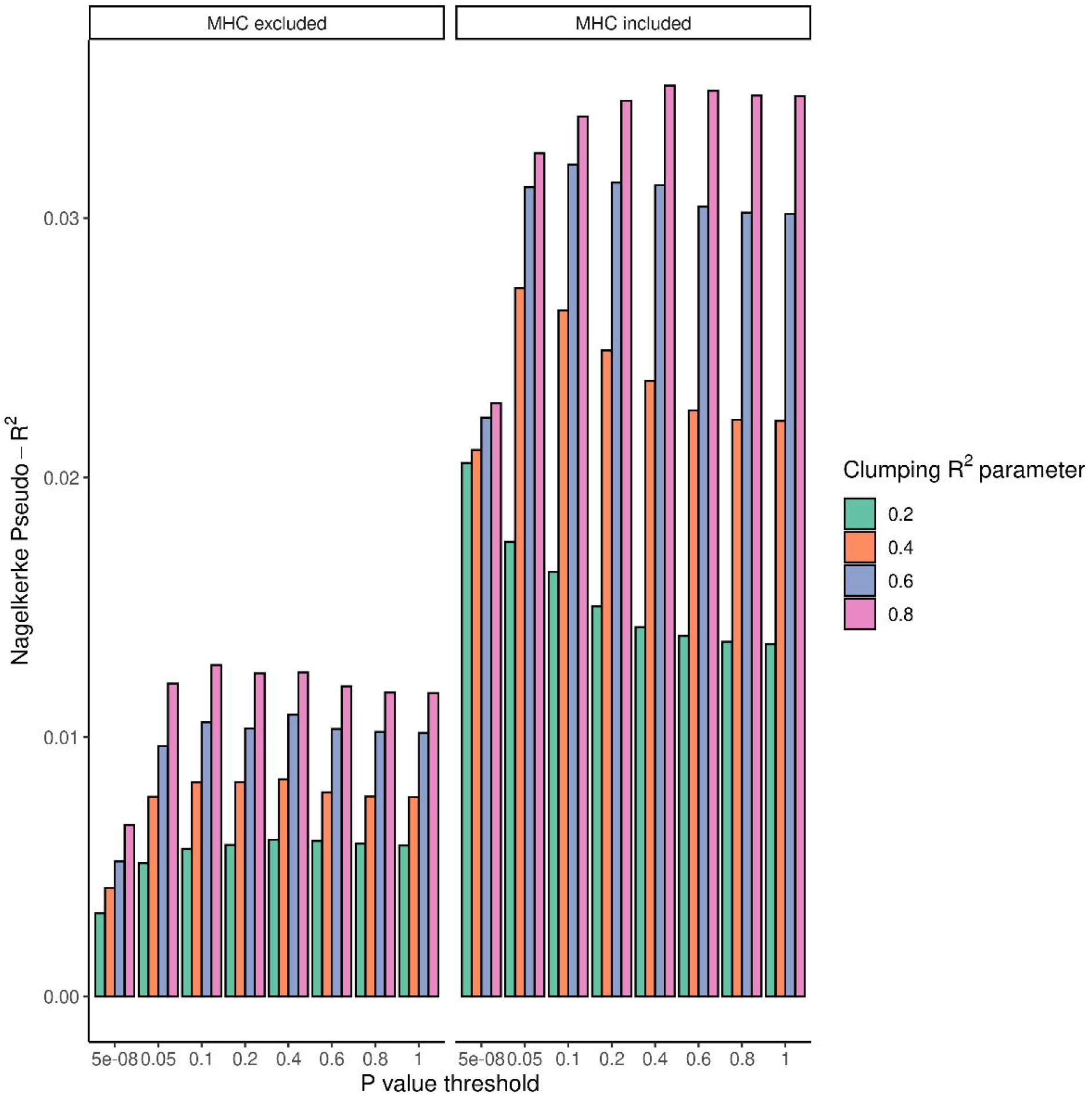
A: Nagelkerke’s pseudo-R^2^ metric for each of the individual PRS used. The R^2^ was calculated by comparing the model fit (age, sex, Townsend deprivation index, the first 4 genetic PCs, and PRS) vs the null model (age, sex, Townsend deprivation index, and the first 4 genetic PCs). A variety of p value thresholds and clumping parameters were used to create different PRS. Note that the clumping R^2^ refers to the linkage disequilibrium threshold within which variants were ‘clumped’, and is a different quantity from the Nagelkerke pseudo-R^2^. PRS are shown both including and excluding the MHC region. B: odds ratios and 95% confidence intervals for MS for individuals in each PRS decile (reference: lowest decile). ORs were calculated from logistic regression models with the following covariates: age, sex, first 4 genetic PCs, and PRS. C: histogram showing PRS distributions among MS cases and controls.

**Figure 3:**
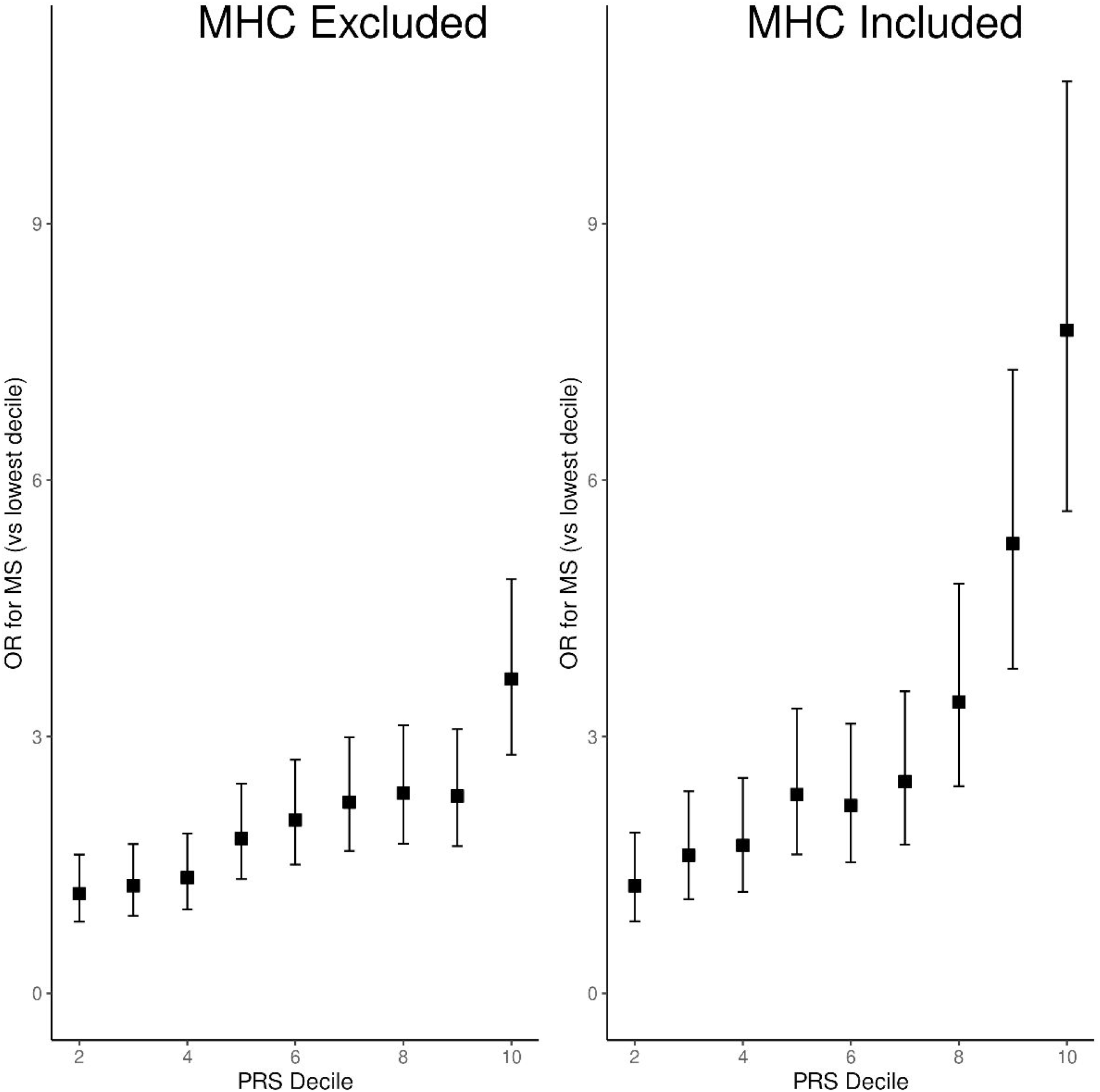
A: Calibration plot showing absolute MS disease probabilities within each PRS decile (of the non-MHC PRS). Other lines represent the mean fitted disease probabilities for models incorporating the MHC PRS, the non-MHC PRS, and null covariates only (age, sex, deprivation, genetic PCs). B: Receiver Operating Characteristic (ROC) curves demonstrating the discriminative performance (i.e. ability to distinguish MS cases from controls) of the null model, MHC PRS, and non-MHC PRS. C&D: scatter plots showing no relationship between PRS (MHC and non-MHC respectively) and normalised age at MS report.

### PRS interactions with environmental risk factors and DRB1*15:01

We found strong evidence of interaction on the additive scale between the PRS_MHC_ and PRS_Non-MHC_ and childhood body size (PRS_MHC_: AP=0.17, 95% CI 0.06 - 0.25, p=0.004; PRS_Non-MHC_: AP=0.17, 95% CI 0.06 - 0.27, p=0.006). We found weaker evidence for interaction on this scale between age at menarche and the PRS_MHC_ (AP=-0.05, 95% CI -0.10 - 0.00, p = 0.033; figure 4a, table 2), but this estimate did not surpass the multiple testing threshold (table 2). There was a lack of strong evidence for other pairwise additive interactions (figure 4) or for multiplicative interactions (supplementary figure 8, supplementary table 6). There was evidence of additive interaction between the PRS_Non-MHC_ and HLA DRB1*15:01 carriage (AP 0.24, 95% CI 0.17 - 0.30, p=0.0002, figure 4b), but no evidence of multiplicative interaction (beta 0.060, p=0.30).

**Table 2:** Attributable Propotion (AP) due to interaction, 95% CIs and two-sided p values for each of the PRS x E interactions examined. CIs represent the 2.5^th^ and 97.5^th^ centile from 10,000 bootstrap replicates. Two-sided p values represent absolute p values with a continuity correction, i.e. for a positive AP, the p value is given as: (number of iterations < 0 + 1)/(total number of iterations+1)*2

| Interaction | AP | Lower CI | Upper CI | P value |
| --- | --- | --- | --- | --- |
| MHC PRS x Childhood body size | 0.167074 | 0.062196 | 0.254741 | 0.0042 |
| Non-MHC PRS x Childhood body size | 0.173705 | 0.055642 | 0.27455 | 0.005599 |
| MHC PRS x Smoking | 0.0768 | -0.05055 | 0.177474 | 0.214179 |
| Non-MHC PRS x Smoking | 0.122975 | -0.00556 | 0.228431 | 0.058794 |
| MHC PRS x Age at menarche | -0.05206 | -0.0968 | -0.00478 | 0.033197 |
| Non-MHC PRS x Age at menarche | 0.021061 | -0.04119 | 0.111064 | 0.551145 |

**Figure 4:**
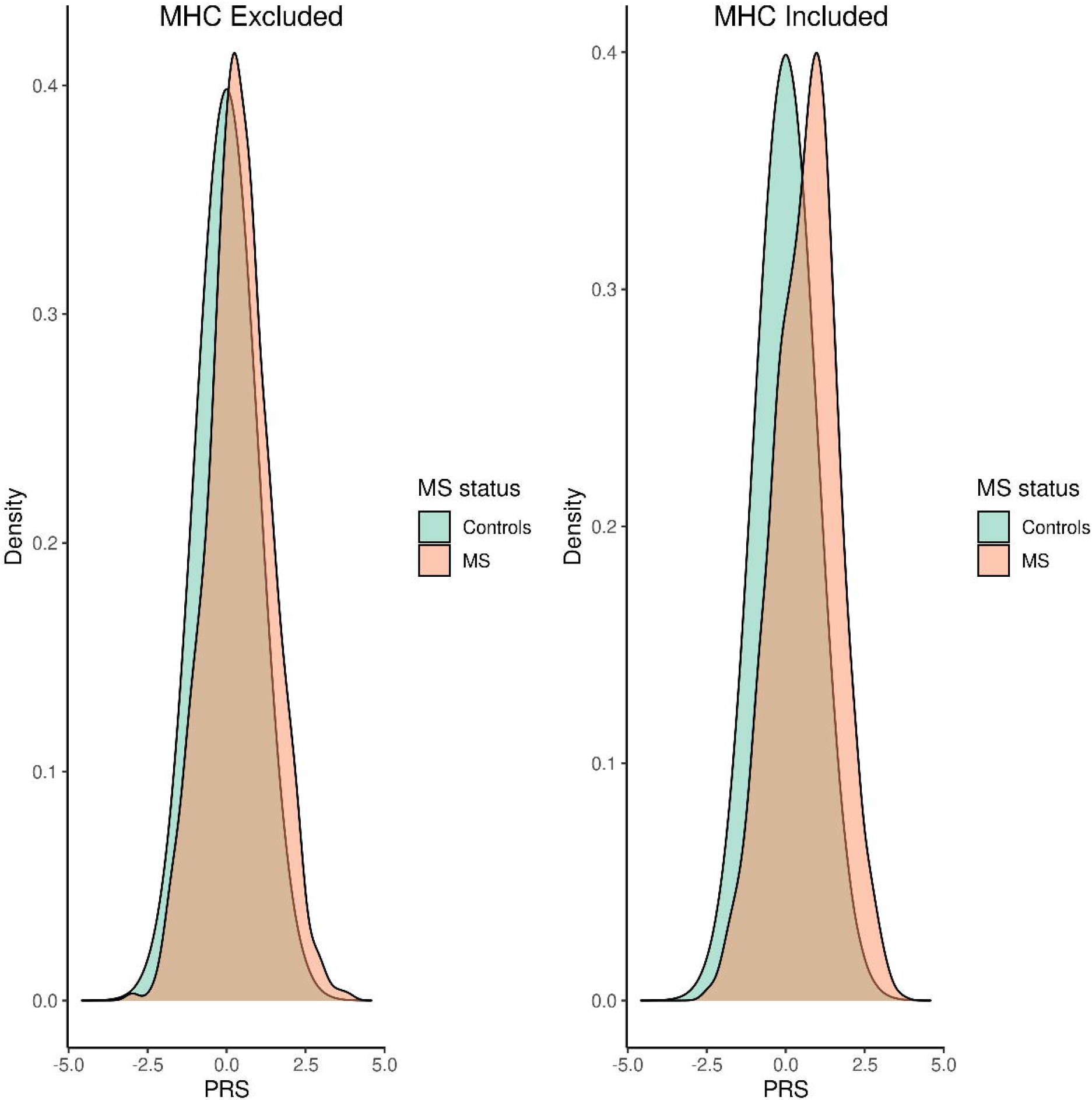
A: Forest plot demonstrating Attributable Proportion due to interaction (AP) and 95% CIs for interactions between environmental exposures and genetic risk factors for MS. If there is no interaction, the AP is 0. AP > 1 indicates positive interaction (combined effects exceed the sum of the individual effects), and vice-versa. CIs are derived from taking the 2.5^th^ and 97.5^th^ percentiles of 10000 bootstrap replicates. B: forest plot demonstrating odds ratios and 95% CIs for participants in the top and bottom PRS deciles, i.e. the highest and lowest 10% of polygenic risk scores. The outcome in each case is MS status, and the exposures of interest are childhood body size, age at menarche, smoking prior to age 20, and carriage of the HLA DRB1*15:01 allele. ORs are from the output of logistic regression model of the form MS risk ∼ Age + Sex + first 4 genetic PCs. Models were built separately for individuals with the highest 10% of genetic risk scores and the lowest 10% of genetic risk scores (‘top’ and ‘bottom’ decile respectively).

## Discussion

In this study we harnessed the scale and breadth of UK Biobank to study >2000 MS cases and >480,000 controls, providing the first evidence that the effect of an established risk factor for MS (childhood obesity) may be potentiated by an individual’s genome-wide genetic risk for MS. We show that this effect persists even when the MHC locus is excluded from the PRS. By using data from the largest GWAS of MS susceptibility to derive and validate polygenic risk scores for MS, both incorporating and excluding the MHC region, we demonstrate supportive evidence for a gene-gene interaction. This work shows that the effect of DRB1*15:01 on MS susceptibility may be potentiated among individuals with a high background genetic risk for MS in this cohort. To our knowledge, our study is the first to demonstrate that the polygenic risk of an individual for MS may alter the effect of established environmental risk factors on their risk of MS^3^. These findings are especially interesting in the context of evidence from mendelian randomisation studies supporting a causal role for childhood obesity in the pathogenesis of MS^21,22^.

Previous studies of gene-environment interactions in MS have focussed on interactions between HLA alleles and environmental risk factors. Specifically, evidence suggests that carriage of high-risk HLA haplotypes containing DRB1*15:01 and lacking A*02:01 enhances the deleterious association of childhood obesity, smoking, Infectious Mononucleosis, and solvent exposure with MS risk^2,789^. The intuitive biological explanation for such interactions is that high-risk HLA alleles may promote presentation of epitopes, e.g. from cigarette smoke or within adipose tissue, in such a way that mimics myelin peptides and triggers CNS-directed autoimmunity. Beyond the MHC, there has been relatively limited study of how genetic variation modulates the effect of environmental risk factors for MS^11,12^, probably in large part due to the relatively small number of datasets with sufficient power, deep phenotyping, and high-quality genetic data required for such analyses.

In this study we created 64 individual PRS, both including and excluding the MHC locus on chromosome 6, which is the strongest single genetic determinant of MS risk and accounts for ∼20% of the SNP heritability of MS in Europeans^1^. Both the non-MHC and MHC PRS were strongly associated with MS risk in both training and testing sets. The non-MHC PRS in this study captured a small proportion of overall MS liability, but was robustly associated with MS. Previous efforts using PRS from the IMSGC explained up to ∼3% of variance^5^. The best-performing non-MHC PRS in this study explained ∼1% of MS variance. This discrepancy could be explained by several factors, including the relatively low number of cases in UK Biobank, the possibility of missed cases, differences in population structure, restriction according to self-declared ethnicity with an additional genetic principal components analysis, and some SNPs not being available and/or failing QC checks in Biobank. Nevertheless, despite low overall variance, the validity of the PRS is underscored by the monotonic relationship between PRS and OR of MS, the robust model fit when using the PRS to model MS risk, reasonable discriminative capacity, and good calibration.

There are several important caveats to this work. Most importantly, while we are able to observe and measure statistical interaction – i.e. deviation from a model whereby the effects of genetic and environmental risk factors are combined additively (in the case of the AP) or multiplicatively (in the case of multiplicative interaction – statistical interaction does not straightforwardly imply biological interaction, nor does it necessarily imply interaction that is meaningful in terms of real-life disease prediction or prevention. We were unable to demonstrate replication in a truly independent cohort (dividing the cohort into training and testing sets does not yield a genuinely independent cohort). Our findings have limited generalisability for non-European groups as UK Biobank participants are predominantly White. MS diagnosis in this cohort are derived from linked healthcare records or self-report, so do not carry the same degree of certainty as criteria-defined MS. Equally, it is conceivable that there are ‘missed’ cases in the dataset, i.e. individuals with MS who do not have a coded diagnosis available through linked healthcare records. However, MS prevalence in UK Biobank approaches the expected UK prevalence, suggesting that the overwhelming majority of individuals with MS are correctly identified. The UKB cohort is highly selected, and is enriched for individuals living near assessment centres, from more affluent socio-economic groups than the general population, for White British individuals, and (intentionally) for individuals over the age of 40 (the minimum age at recruitment). These factors carry a risk of introducing various biases, e.g. through collider bias, which may induce spurious associations and destroy true associations. Our findings concerning gene-gene interactions could be replicated in ‘genetics-only’ cohorts such as the IMSGC, and we would encourage others to attempt to replicate this finding in large GWAS cohorts (with many more cases than the ∼2000 in UKB) so we can ascertain whether it is robust.

The key variables used in this study are retrospective or cross-sectional (e.g. MS diagnosis, self-reported body size in childhood, self-reported smoking status). Not only are these subject to recall bias, but more importantly our results are not revealing about predicting an individual’s risk of developing MS. To demonstrate *predictive* power, these results need to be replicated in a longitudinal cohort. In addition, the metric we focus on, ‘comparative body size at age 10’, is clearly not a perfect proxy for childhood obesity. Other limitations to this study include the limited overall variance explained by both optimal PRS, the relatively small absolute number of people with MS, and the imperfect nature of self-reported phenotypes. Furthermore, some exposures known to be strongly associated with MS were either unavailable (e.g. vitamin D status prior to diagnosis) or so under-reported as to be unreliable (e.g. Infectious Mononucleosis).

Despite these limitations, our study also has some strengths. We use the UK Biobank dataset, which provides a unique opportunity to study gene-environment interactions on a large-scale. The vast number of controls in UKB adds substantial power. We tune and test the PRS in separate samples, which is important to prevent overfitting of the PRS to the data. We use an agnostic approach to develop the PRS, using a range of clumping-and-thresholding parameters to discover the optimal structure of the PRS, allowing us to discover a significant improvement in predictive power from using a large number of variants weakly associated with MS over using strictly ‘GWAS-significant’ hits (p<5e-8). These optimal parameters also reiterate the polygenic architecture of MS.

We evaluate interactions on both the multiplicative and additive scales, as has become standard practice to avoid missing biologically-significant interactions^2^. We additionally evaluate the relationship between the PRS and proxies for clinical characteristics of MS, including age at diagnosis and disability status. We evaluated whether the effect of DRB1*15:01 is modulated by polygenic risk, as has been demonstrated for high-effect variants in the LDL-R (causing Familial Hypercholesterolaemia) and BRCA (causing breast cancer)^23^, and find evidence in support of this hypothesis. Clearly, this finding is easily replicated in the IMSGC cohort and we would urge caution in overinterpreting the finding without confirmation in this far larger cohort of cases.

This study thus provides novel evidence that childhood body size interacts with non-HLA MS genetic risk. Demonstrating benefit for preventive measures in rare, complex diseases like MS is a challenge due to the low population incidence and the small effects of individual interventions. Power can be enhanced by enriching for high-risk individuals, and by selecting individuals who are likely to experience the greatest benefit from the intervention. As the effect of childhood body size on MS risk appears greater among individuals with a high genome-wide genetic risk, trials attempting to demonstrate benefit of targeting childhood obesity may benefit from risk-stratifying individuals using this approach. Further efforts are required to localise the variants and genes which account for the observed interaction effects, which should help to shed further light on the biology of these risk factors and improve efforts to individualise MS risk prediction algorithms in the future.

## Acknowledgements and data availability

We would like to thank the relevant consortia for making their data available. MS GWAS data were taken from the MS Chip discovery summary statistics. IMSGC summary statistics are available via request on the website: https://nettskjema.no/answer/imsgc-data-access.html. We would like to thank the Queen Mary University High Performance Computing team for their help with computing resources. We would like to thank the participants and researchers involved in UK Biobank, who have created an exceptional resource. UK Biobank data are available on request through their website. Code used in this paper is available on Github (@benjacobs123456).

---

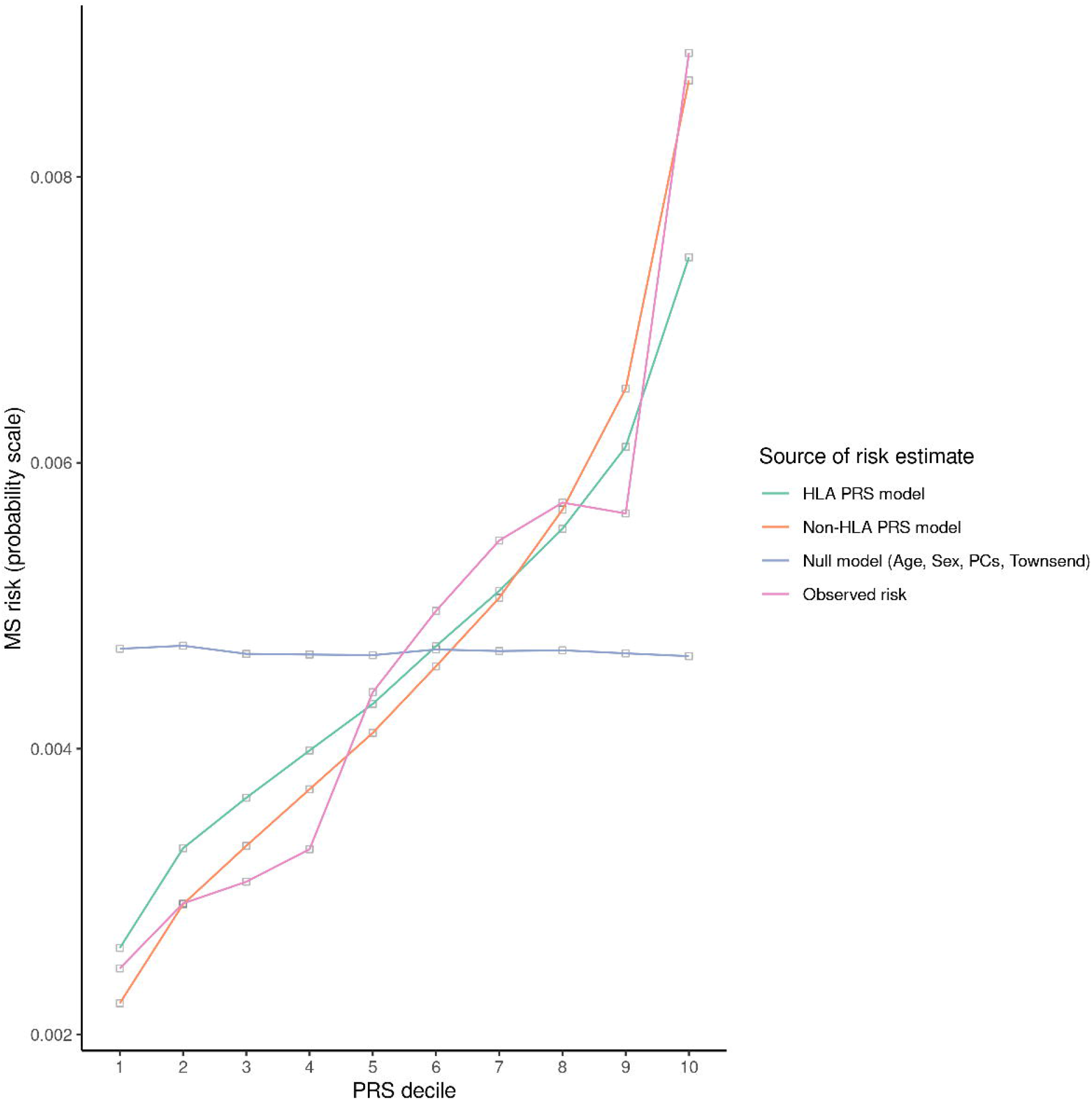

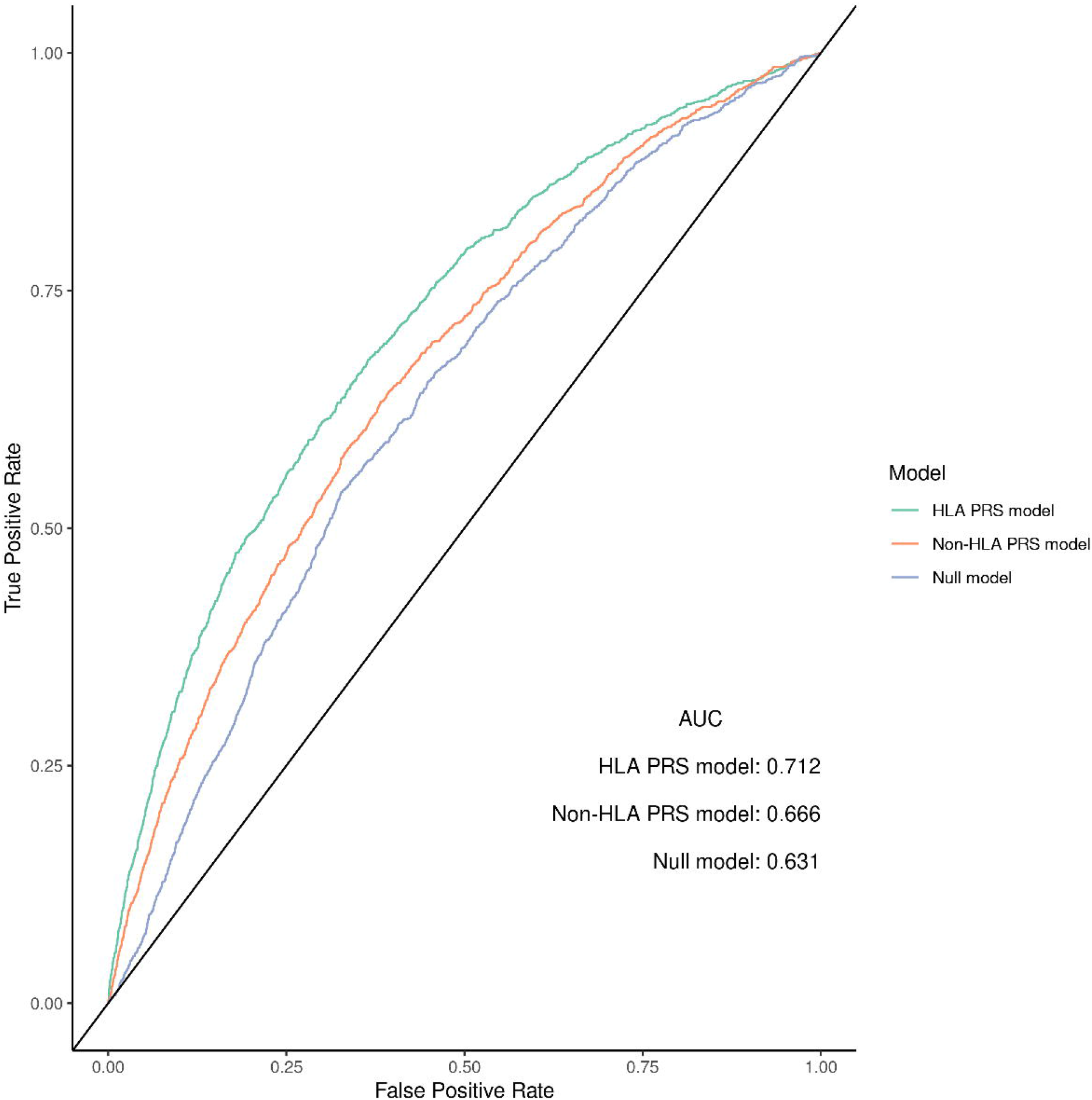

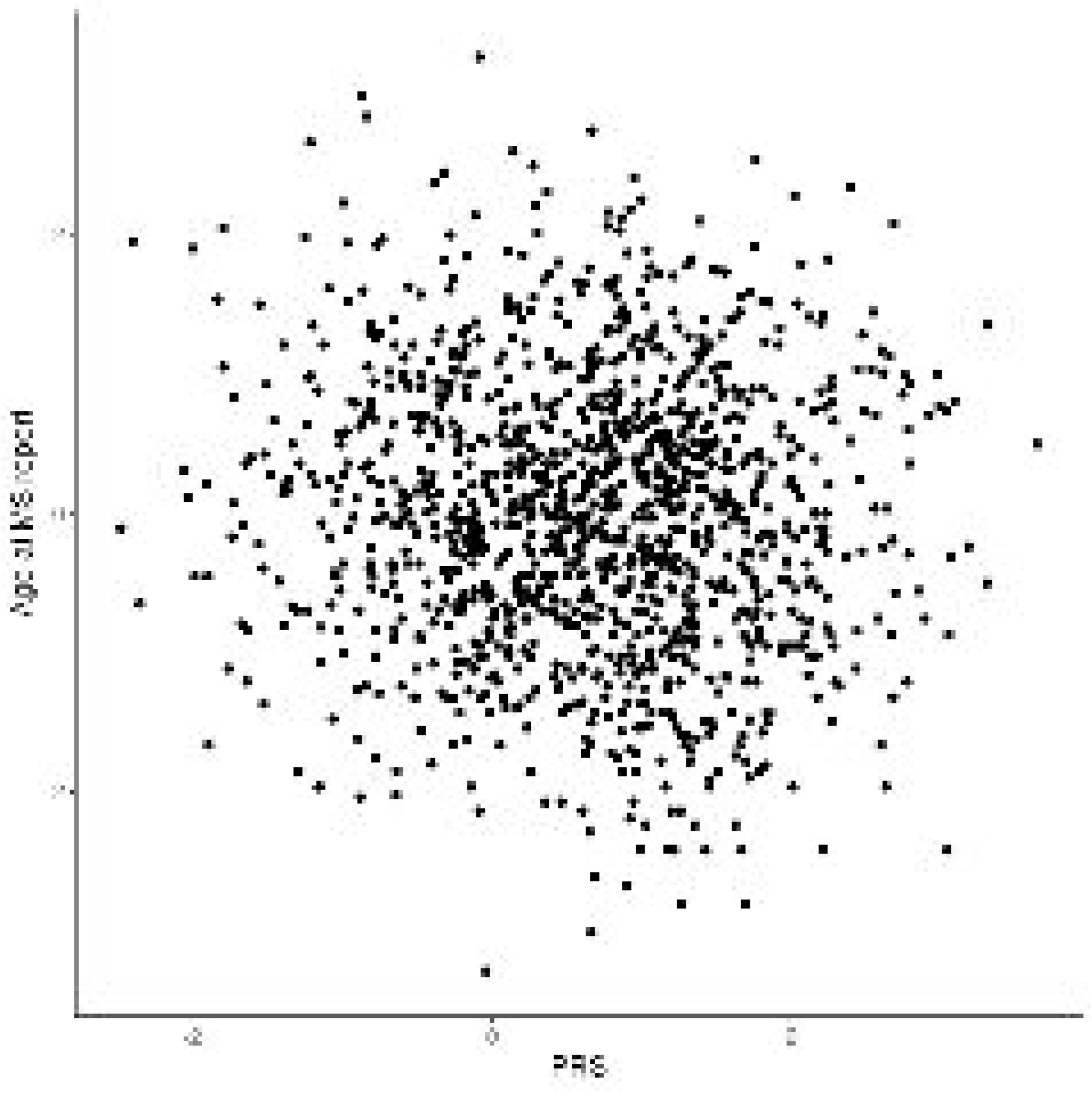

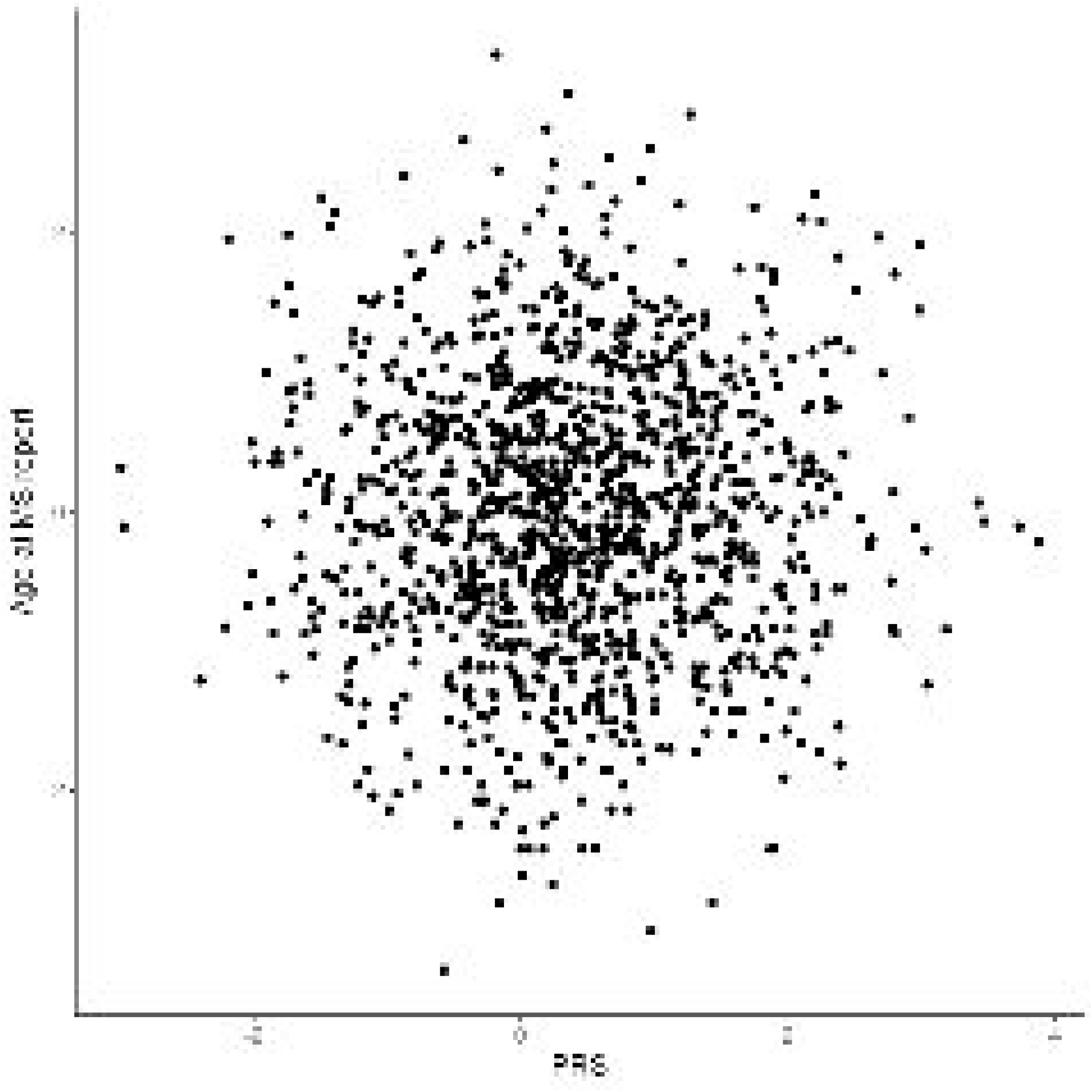

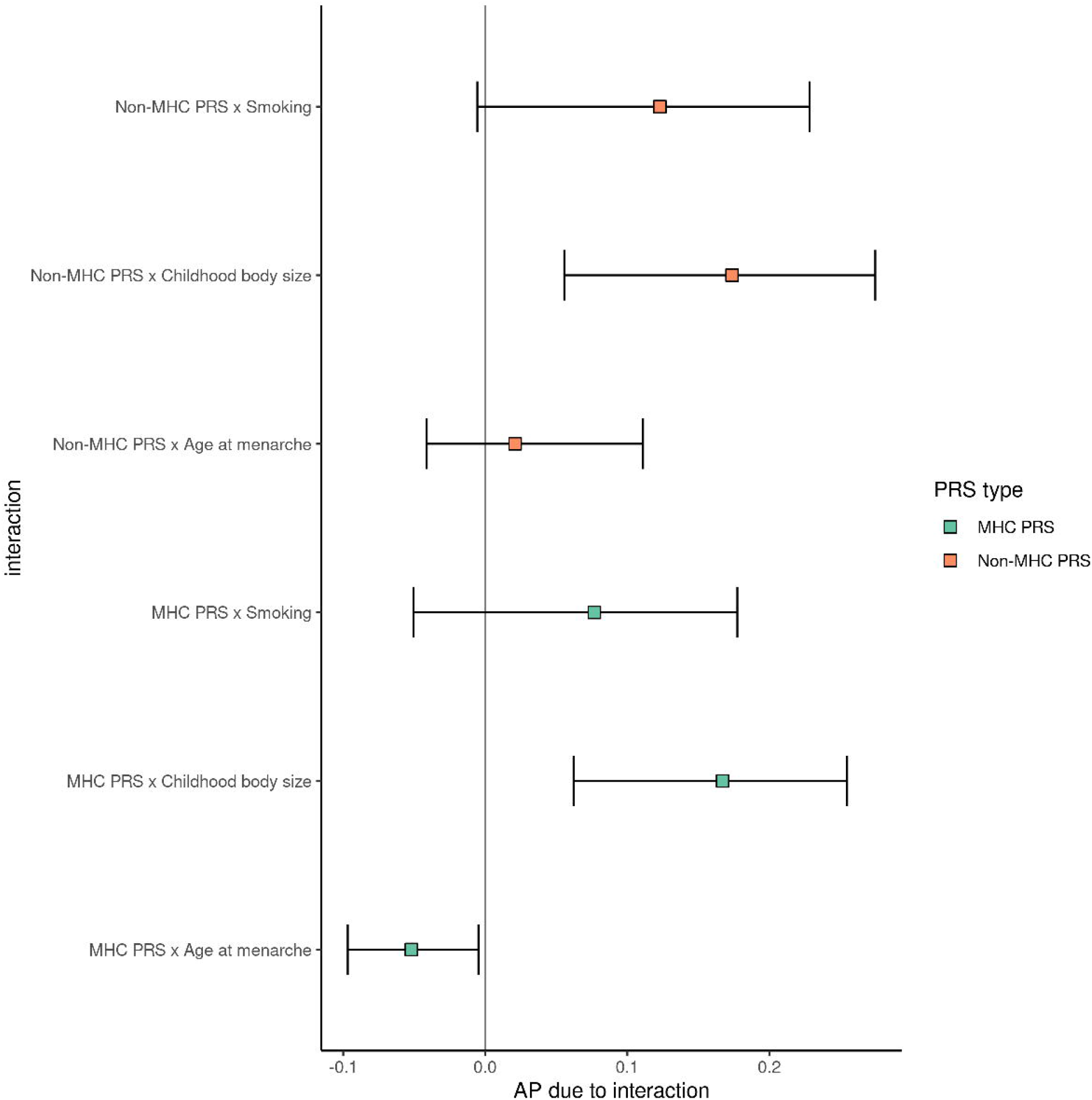

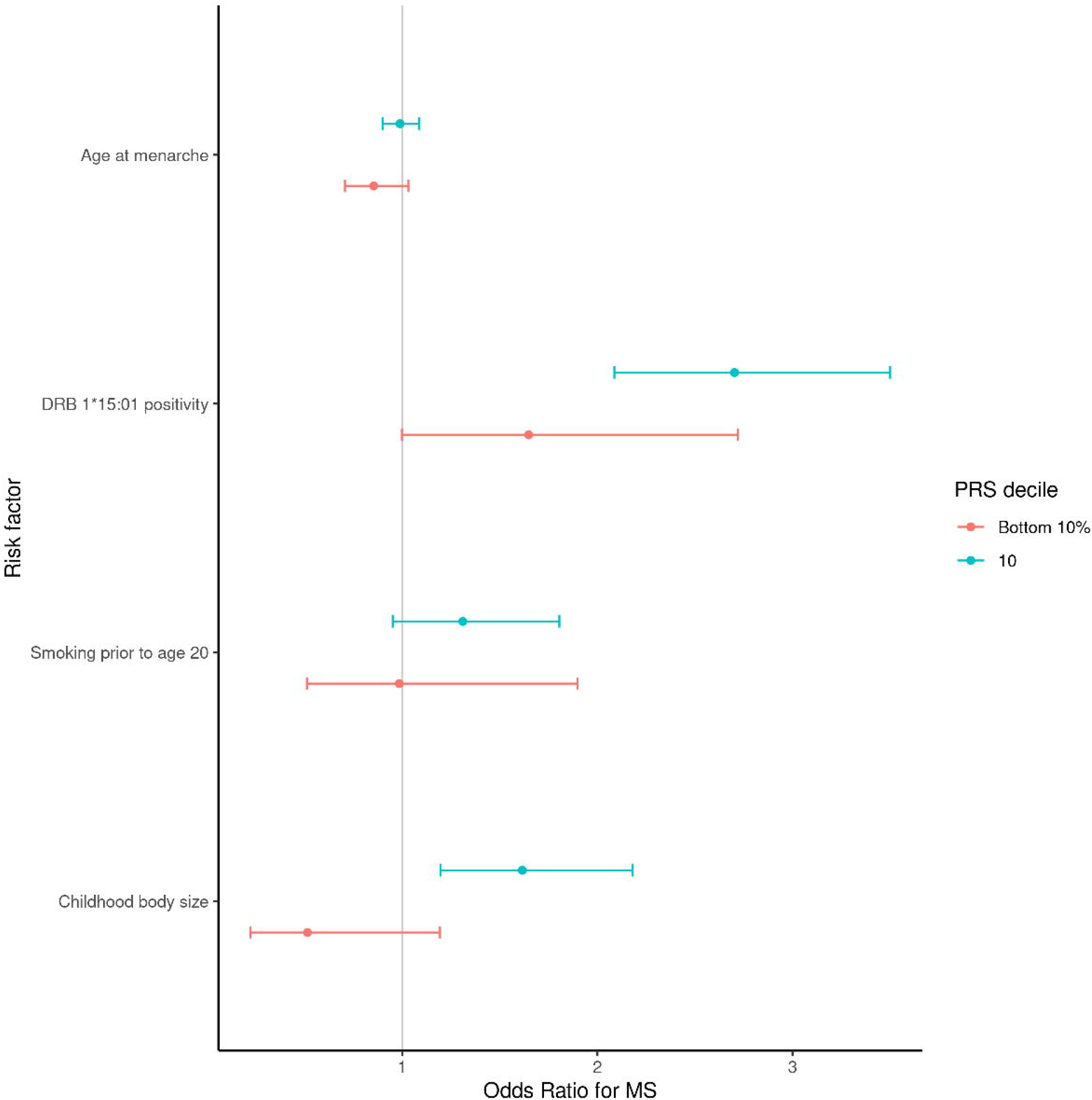

