## supplemental data for "Gene-environment interactions in Multiple Sclerosis: a UK Biobank study"

Supplementary methods

*Definition of cases and controls, definition of age at disease onset*

ICD codes in UK Biobank are extracted from Hospital Episode Statistics and refer to diagnoses recorded during a hospital admission/encounter. We included all participants with any MS code as determined by the UK Biobank field ‘Source of report of G35 (multiple sclerosis)’ (see supplementary table 1 for details of all exposure/outcome definitions). Overall 2365 participants with MS were present in the dataset. Genetic data were available for 2276 MS cases and 486,000 unmatched controls. The breakdown of diagnostic codes is shown in supplementary table 2. Age at diagnosis was determined using the date of the first recorded MS diagnostic code (using UK Biobank field ‘date G35 first reported’). MS diagnoses reported before age 20 were excluded leaving 2250 participants in the main case-control analysis.

*Polygenic risk score derivation*

The following steps were applied in order to generate 64 individual risk scores:

1. We extracted variant associations with MS from the discovery stage meta-analysis summary statistics obtained from the IMSGC^1^.
2. We excluded sex-chromosome variants, variants with strand-ambiguous alleles (A/T and C/G SNPs), variants without an rsid, duplicate variants, variants with mismatched alleles between datasets, variants not typed/imputed/sequenced in all 3 datasets (reference 1000 genomes samples, IMSGC meta-analysis summary statistics, and UK Biobank), variants with high missingness (>10%), deviation from Hardy-Weinberg equilibrium (p<1e-5), variants with low imputation quality (Mach R^2^ < 0.3), and variants with MAF < 0.01 in either UK Biobank or 1000 genomes European samples.
3. We created scores either including or excluding the MHC region. For scores excluding the MHC region (‘non-MHC PRS’), we excluded SNPs within the extended MHC (chr6:25,000,000 to chr6:35,000,000 on hg19).
4. We included variants with association statistic p values (for association with MS, obtained from IMSGC meta-analysis summary statistics) below a series of  arbitrary p value thresholds (5e-8, 0.05, 0.1, 0.2, 0.4, 0.6, 0.8, and 1).
5. We clumped using several r^2^ thresholds (0.2, 0.4, 0.6, 0.8) and a clumping distance of 250kBP. Reference genome data were obtained from the 503 participants of European ancestry in the 1000 genomes project^16^.

This procedure was used to generate 64 different computations of PRS (32 including MHC and 32 MHC-excluded scores). Beta coefficients (i.e. log Odds Ratios) were derived from the IMSGC discovery GWAS^1^. We applied these 64 PRS to individuals in UK Biobank after applying rigorous individual quality control (QC): individuals with >10% missing genotypes were excluded, and only individuals with self-reported ‘British’ ethnicity and genetic ‘White European’ ancestry as defined by genetic principal components were included (supplementary figures 3 & 4 show the degree of population stratification prior to removal of PCA outliers and how this is mitigated by restricting to European participants). One of each pair of related individuals (kinship coefficient > 0.0844) was excluded to reduce the risk of cryptic relatedness biasing the results (supplementary figure 1). Effect allele dosage at each locus was multiplied by the beta coefficient (log OR) to generate the risk score for that locus. Scores were standardised to have mean 0 and unit variance (i.e. variance = 1) for each SNP. Where genotypes were missing, the score at that locus was defined as the mean of all scores at that locus. Risk scores were totalled across the genome to calculate a score for each individual. Derivation of individual risk scores was implemented in PLINK2 using the ‘--score’ flag^17^.

*Polygenic risk score validation in the training set (70% of the cohort)*

In addition to determining the variance explained in MS risk by the optimal PRS, we performed further validation tests. We calculated odds ratios for MS for each decile of the PRS compared to the lowest decile using multivariable logistic regression models controlling for age, sex, Townsend score, and the first four genetic PCs. Subsequently, for each PRS decile, we calculated the observed absolute risk of MS (i.e. number of cases / total number of participants in decile). We estimated probabilities of MS from either a full model (PRS, age, sex, Townsend, first 4 PCs) or a null model (age, sex, Townsend, first 4 PCs) by building logistic regression models with these covariates and obtaining the fitted probabilities for each individual. We averaged these fitted probabilities within each decile to give an estimate of the mean fitted MS probability under each model, and to allow comparison with the observed risk. Finally, we calculated Receiver Operating Characteristic (ROC) scores to quantify the classified performance of the PRS. To adjust for confounding, we used the fitted probabilities from the above models as input data for ROC analysis. We included a null model (with the same covariates as above) for comparison.

*Additive and multiplicative interaction*

Interaction was assessed on both the additive and multiplicative scales. Interaction on the additive scale was assessed by calculating the Attributable Proportion due to interaction (AP). Additive interaction analyses were based on multivariable logistic regression models incorporating age, sex, and the first four genetic principal components as confounders^18^.

For a logistic regression model of the form:

$$log\frac{p}{1-p}= \beta_{0}+\beta_{1}x+\beta_{2}y+\beta_{3}xy$$

In which *log(p/1-p)* is the log odds of MS, *x* and *y* are the values of exposure variables (e.g. childhood body size, smoking, polygenic risk score), and $xy$ is the interaction term, then the Relative Excess Risk due to Interaction (RERI) can be calculated as:

$$RERI=\exp\left( \beta_{1}+ \beta_{2}+\beta_{3} \right)-\exp\left( \beta_{1} \right)-\exp\left( \beta_{2} \right)+1$$

The AP can be conceived of as the proportion of the disease in the doubly-exposed group attributable to the interaction between the risk factors, i.e:

$$AP= \frac{RERI}{\exp\left( \beta_{1}+ \beta_{2}+\beta_{3} \right)}$$

This model can be expanded to include confounding covariates, in which case the beta coefficients are adjusted for confounders^18^. We restricted this analysis to participants with genetically European ancestry determined by both self-report and genetic ethnic grouping as above.

For interaction analyses, covariates were age, sex, and the first four genetic principal components. The PRS was transformed using the inverse-normal transformation and treated as a continuous variable. Confidence intervals for the AP were estimated using bootstrap resampling of the entire dataset with replacement for 10,000 iterations^18^ with 95% confidence intervals derived from the 2.5th and 97.5th centile values. Two-sided exact P values were calculated with a continuity correction as number of iterations more extreme than the observed AP + 1 / total number of iterations + 1 * 2:

$$p=2 \times\frac{1+ number of iterations<0}{10001}$$

Interaction on the multiplicative scale was assessed using a logistic regression model incorporating an interaction term and quantified using the likelihood ratio.

Supplementary table 1: definitions of covariates extracted from UK Biobank, including links to the relevant fields in the UK Biobank data showcase.

| **Variable** | **Method of data acquisition** | **UK Biobank URL** |
| --- | --- | --- |
| Sex | Registry | <http://biobank.ndph.ox.ac.uk/showcase/field.cgi?id=31> |
| Month of birth | Registry | <http://biobank.ndph.ox.ac.uk/showcase/field.cgi?id=52> |
| Age at recruitment | Derived from birth date and date of 1st assessment | <http://biobank.ndph.ox.ac.uk/showcase/field.cgi?id=21022> |
| Place of Birth in UK North Co-ordinate (latitude) | Verbal interview | <http://biobank.ndph.ox.ac.uk/showcase/field.cgi?id=129> |
| Townsend Deprivation  Index at recruitment | Derived from census data and postcode | <http://biobank.ndph.ox.ac.uk/showcase/field.cgi?id=189> |
| Breastfeeding status | Touchscreen question: “Were you breastfed when you were a baby?” | <http://biobank.ndph.ox.ac.uk/showcase/field.cgi?id=1677> |
| Age at voice breaking | Touchscreen question "When did your voice break?" | <http://biobank.ndph.ox.ac.uk/showcase/field.cgi?id=2385> |
| Age at menarche | Touchscreen question "How old were you when your periods started?" | <http://biobank.ndph.ox.ac.uk/showcase/field.cgi?id=2714> |
| Age first had sexual intercourse | Touchscreen question "What was your age when you first had sexual intercourse? (Sexual intercourse includes vaginal, oral or anal intercourse)" | <http://biobank.ndph.ox.ac.uk/showcase/field.cgi?id=2139> |
| Smoking status | Derived from 2 touchscreen questions:   1. "Do you smoke tobacco now?" 2. "In the past, how often have you smoked tobacco?"   And for current/previous smokers, age when started smoking was derived from the question "How old were you when you first started smoking on most days?" | <http://biobank.ndph.ox.ac.uk/showcase/field.cgi?id=20116>  <http://biobank.ndph.ox.ac.uk/showcase/field.cgi?id=3436>  <http://biobank.ndph.ox.ac.uk/showcase/field.cgi?id=2867> |
| Comparative body size aged 10 | Touchscreen question: "When you were 10 years old, compared to average would you describe yourself as:" | <http://biobank.ndph.ox.ac.uk/showcase/field.cgi?id=1687> |
| Ethnicity | Derived from several touchscreen questions | <http://biobank.ndph.ox.ac.uk/showcase/field.cgi?id=21000> |
| Birth weight (Kg) | Touchscreen | <http://biobank.ndph.ox.ac.uk/showcase/field.cgi?id=20022> |
| Infectious mononucleosis | Self-reported | <http://biobank.ndph.ox.ac.uk/showcase/field.cgi?id=20002> |
| Multiple sclerosis | HES records -  ICD10 G35; ICD9 3409  Self-reported  Death register  GP records | http://biobank.ndph.ox.ac.uk/showcase/field.cgi?id=131043 |
| Maternal smoking around the time of birth | Touchscreen question: "Did your mother smoke regularly around the time when you were born?" | <http://biobank.ndph.ox.ac.uk/showcase/field.cgi?id=1787> |
| Other diseases (PD, SLE, RA, IBD etc) | HES records (ICD codes) | http://biobank.ctsu.ox.ac.uk/crystal/field.cgi?id=41270 |

Supplementary table 2: source of MS report (derived from UK Biobank source of first G35 report).

| **Source of MS report** | **N** |
| --- | --- |
| death register only | 2 |
| HES data and other source/s | 100 |
| HES data only | 272 |
| primary care and other source/s | 437 |
| primary care only | 137 |
| self-report and other source/s | 1118 |
| self-report only | 299 |

Supplementary table 3: Predictors which conferred a good model fit (likelihood ratio p value < multiple testing threshold for alpha=0.05) from the case-control study were combined in multivariable model adjusting for HLA DRB1*15:01 and A*02:01 alleles. For menarche, sex was not included as a covariate as it applies to women only.

| **Exposure** | **OR (95% CI)** | **P Value** |
| --- | --- | --- |
| Comparative body size aged 10: about average | 1.19 (1.08 to 1.32) | 0.000712 |
| Comparative body size aged 10: plumper | 1.36 (1.2 to 1.55) | 1.95E-06 |
| Smoking status prior to age 20 | 1.19 (1.07 to 1.33) | 0.001582 |
| DRB1*15:01: 1 | 2.51 (2.29 to 2.75) | 1.45E-87 |
| DRB1*15:01: 2 | 4.98 (4.17 to 5.94) | 2.14E-71 |
| A*02:01: 1 | 0.67 (0.61 to 0.73) | 9.49E-17 |
| A*02:01: 2 | 0.58 (0.47 to 0.71) | 6.76E-08 |
| Age at menarche | 0.95 (0.92 to 0.98) | 0.003745 |

Supplementary table 4: Nagelkerke’s Pseudo-R2 metrics for each of the 64 PRS created. Column 1 indicated the P value threshold for variants included in the score. Column 2 indicated the clumping R2 metric used to define independent variants for the score. Column 3 indicates the Nagelkerke Pseudo-R2 metric, quantifying the proportion of total MS risk explained by the PRS in the training set. Column 4 indicates whether the score included or excluded variants within the super-extended MHC.

| P value threshold | Clumping R2 threshold | Nagelkerke's Pseudo R2 | MHC or non-MHC score |
| --- | --- | --- | --- |
| 5E-08 | 0.8 | 0.00660754 | MHC excluded |
| 5E-08 | 0.6 | 0.00520834 | MHC excluded |
| 5E-08 | 0.4 | 0.00418614 | MHC excluded |
| 5E-08 | 0.2 | 0.00321115 | MHC excluded |
| 0.05 | 0.8 | 0.0120634 | MHC excluded |
| 0.05 | 0.6 | 0.00965529 | MHC excluded |
| 0.05 | 0.4 | 0.00770067 | MHC excluded |
| 0.05 | 0.2 | 0.00514216 | MHC excluded |
| 0.1 | 0.8 | 0.0127784 | MHC excluded |
| 0.1 | 0.6 | 0.0105706 | MHC excluded |
| 0.1 | 0.4 | 0.00825061 | MHC excluded |
| 0.1 | 0.2 | 0.00569735 | MHC excluded |
| 0.2 | 0.8 | 0.0124595 | MHC excluded |
| 0.2 | 0.6 | 0.0103313 | MHC excluded |
| 0.2 | 0.4 | 0.00825553 | MHC excluded |
| 0.2 | 0.2 | 0.00584438 | MHC excluded |
| 0.4 | 0.8 | 0.0124931 | MHC excluded |
| 0.4 | 0.6 | 0.0108599 | MHC excluded |
| 0.4 | 0.4 | 0.00837708 | MHC excluded |
| 0.4 | 0.2 | 0.00604783 | MHC excluded |
| 0.6 | 0.8 | 0.0119561 | MHC excluded |
| 0.6 | 0.6 | 0.0103136 | MHC excluded |
| 0.6 | 0.4 | 0.00787574 | MHC excluded |
| 0.6 | 0.2 | 0.00599818 | MHC excluded |
| 0.8 | 0.8 | 0.0117144 | MHC excluded |
| 0.8 | 0.6 | 0.0102012 | MHC excluded |
| 0.8 | 0.4 | 0.00770732 | MHC excluded |
| 0.8 | 0.2 | 0.00589313 | MHC excluded |
| 1 | 0.8 | 0.0117006 | MHC excluded |
| 1 | 0.6 | 0.0101663 | MHC excluded |
| 1 | 0.4 | 0.00768628 | MHC excluded |
| 1 | 0.2 | 0.00582148 | MHC excluded |
| 5E-08 | 0.8 | 0.0228652 | MHC included |
| 5E-08 | 0.6 | 0.0223136 | MHC included |
| 5E-08 | 0.4 | 0.0210648 | MHC included |
| 5E-08 | 0.2 | 0.0205637 | MHC included |
| 0.05 | 0.8 | 0.0325017 | MHC included |
| 0.05 | 0.6 | 0.0311873 | MHC included |
| 0.05 | 0.4 | 0.0272963 | MHC included |
| 0.05 | 0.2 | 0.0175248 | MHC included |
| 0.1 | 0.8 | 0.0339157 | MHC included |
| 0.1 | 0.6 | 0.0320654 | MHC included |
| 0.1 | 0.4 | 0.0264375 | MHC included |
| 0.1 | 0.2 | 0.0163644 | MHC included |
| 0.2 | 0.8 | 0.0345205 | MHC included |
| 0.2 | 0.6 | 0.0313649 | MHC included |
| 0.2 | 0.4 | 0.0248961 | MHC included |
| 0.2 | 0.2 | 0.0150374 | MHC included |
| 0.4 | 0.8 | 0.0351071 | MHC included |
| 0.4 | 0.6 | 0.0312699 | MHC included |
| 0.4 | 0.4 | 0.0237235 | MHC included |
| 0.4 | 0.2 | 0.0142392 | MHC included |
| 0.6 | 0.8 | 0.0349039 | MHC included |
| 0.6 | 0.6 | 0.0304377 | MHC included |
| 0.6 | 0.4 | 0.0225894 | MHC included |
| 0.6 | 0.2 | 0.0138951 | MHC included |
| 0.8 | 0.8 | 0.034721 | MHC included |
| 0.8 | 0.6 | 0.03021 | MHC included |
| 0.8 | 0.4 | 0.0222316 | MHC included |
| 0.8 | 0.2 | 0.0136718 | MHC included |
| 1 | 0.8 | 0.0346988 | MHC included |
| 1 | 0.6 | 0.0301573 | MHC included |
| 1 | 0.4 | 0.0221831 | MHC included |
| 1 | 0.2 | 0.0135759 | MHC included |

Supplementary table 5: odds ratios for MS in each PRS decile vs the lowest decile. Multivariable logistic regression models were built separately for the MHC and non-MHC PRS scores. Data are from the testing (validation) set of participants.

| **MHC PRS** |  |  |  |  |
| --- | --- | --- | --- | --- |
| PRS decile | OR | Lower CI | Upper CI | P |
| 2 | 1.256948 | 0.841679 | 1.877103 | 0.263716 |
| 3 | 1.612257 | 1.10115 | 2.360597 | 0.014076 |
| 4 | 1.729441 | 1.187062 | 2.51964 | 0.004329 |
| 5 | 2.323872 | 1.623769 | 3.325833 | 4.02E-06 |
| 6 | 2.195242 | 1.529516 | 3.150727 | 2.00E-05 |
| 7 | 2.474792 | 1.735465 | 3.52908 | 5.60E-07 |
| 8 | 3.405705 | 2.421977 | 4.788993 | 1.84E-12 |
| 9 | 5.2579 | 3.791414 | 7.291609 | 2.56E-23 |
| 10 | 7.752433 | 5.637506 | 10.66078 | 2.10E-36 |
| **Non-MHC PRS** |  |  |  |  |
| PRS decile | OR | Lower CI | Upper CI | P |
| 2 | 1.164714 | 0.836001 | 1.622676 | 0.36746 |
| 3 | 1.257403 | 0.906804 | 1.743555 | 0.169627 |
| 4 | 1.352929 | 0.980615 | 1.866599 | 0.065652 |
| 5 | 1.808435 | 1.334063 | 2.451486 | 1.35E-04 |
| 6 | 2.025946 | 1.503953 | 2.729113 | 3.41E-06 |
| 7 | 2.232766 | 1.664918 | 2.994287 | 8.11E-08 |
| 8 | 2.340004 | 1.748563 | 3.131497 | 1.07E-08 |
| 9 | 2.304899 | 1.720787 | 3.087286 | 2.14E-08 |
| 10 | 3.674777 | 2.789896 | 4.84032 | 2.05E-20 |

Supplementary table 6: Interaction terms for interaction between PRS and environmental exposures. P values represent Wald test P values (i.e. testing the null hypothesis that the interaction term beta = 0).

| Interaction | Beta | SE | P |
| --- | --- | --- | --- |
| MHC PRS x Childhood body size | 0.131579 | 0.072842 | 0.070862569 |
| Non-MHC PRS x Childhood body size | 0.152394 | 0.071829 | 0.033869511 |
| MHC PRS x Smoking | 0.016847 | 0.073197 | 0.817965885 |
| Non-MHC PRS x Smoking | 0.099224 | 0.072081 | 0.168646678 |
| MHC PRS x Age at menarche | -0.0256 | 0.021464 | 0.233002052 |
| Non-MHC PRS x Age at menarche | 0.026065 | 0.021268 | 0.220362814 |
