## supplemental figures for "Gene-environment interactions in Multiple Sclerosis: a UK Biobank study"

Supplementary figure 1: participant flow through the study. The number of individuals at each stage used for analysis is highlighted in the boxes with black outline. The boxes with grey outline indicate the filtering criterion/criteria used at each stage.

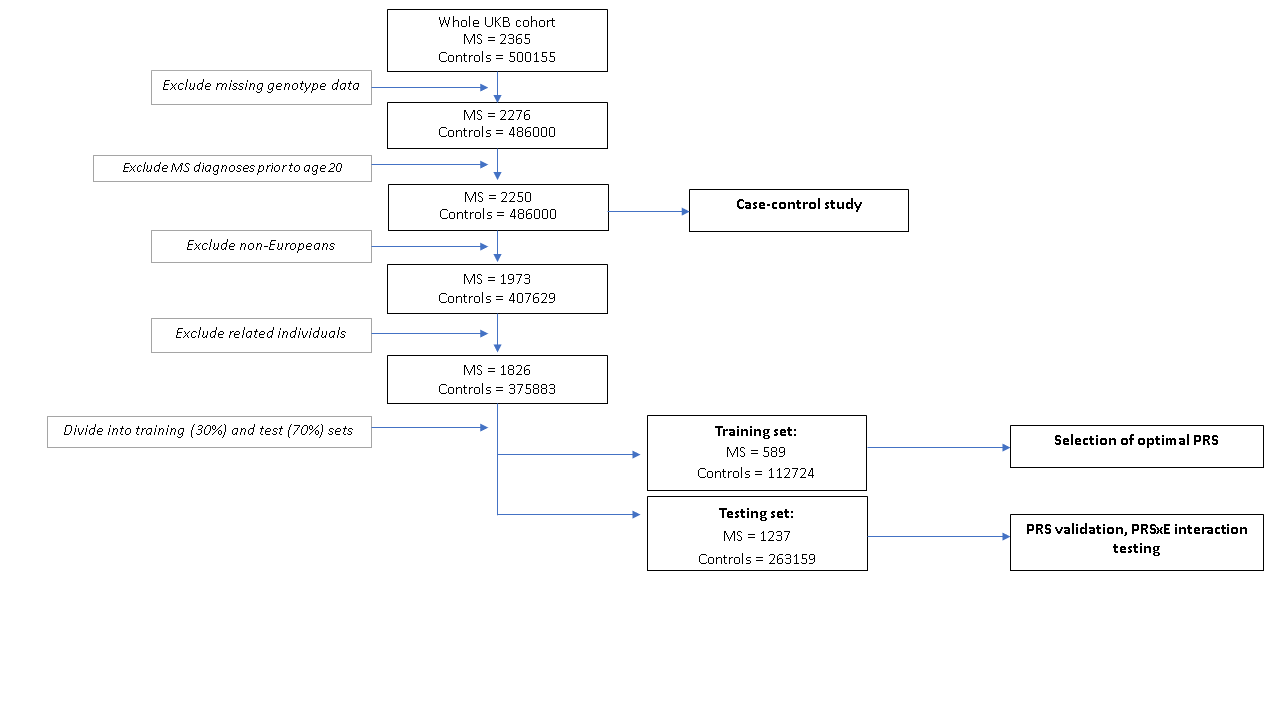

Supplementary figure 2: density plot depicting age at MS diagnostic code report. Note that these codes refer to the first reported diagnosis, not the first symptom onset, and are at best a proxy for disease onset.

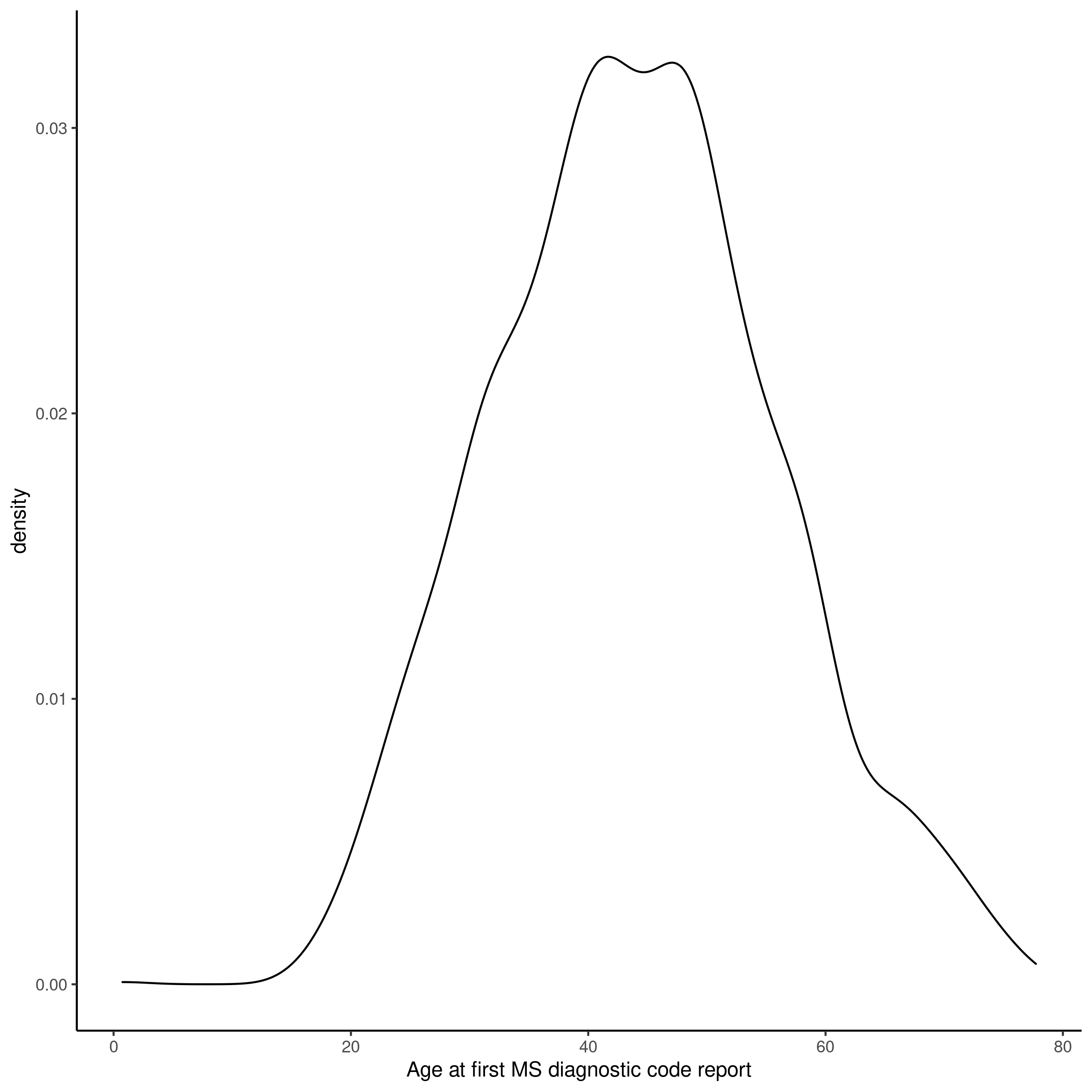

Supplementary figure 3: Genetic principal component plot depicting PCs 1 and 2 prior to ancestry exclusions. Self-reported ethnicity is shown.
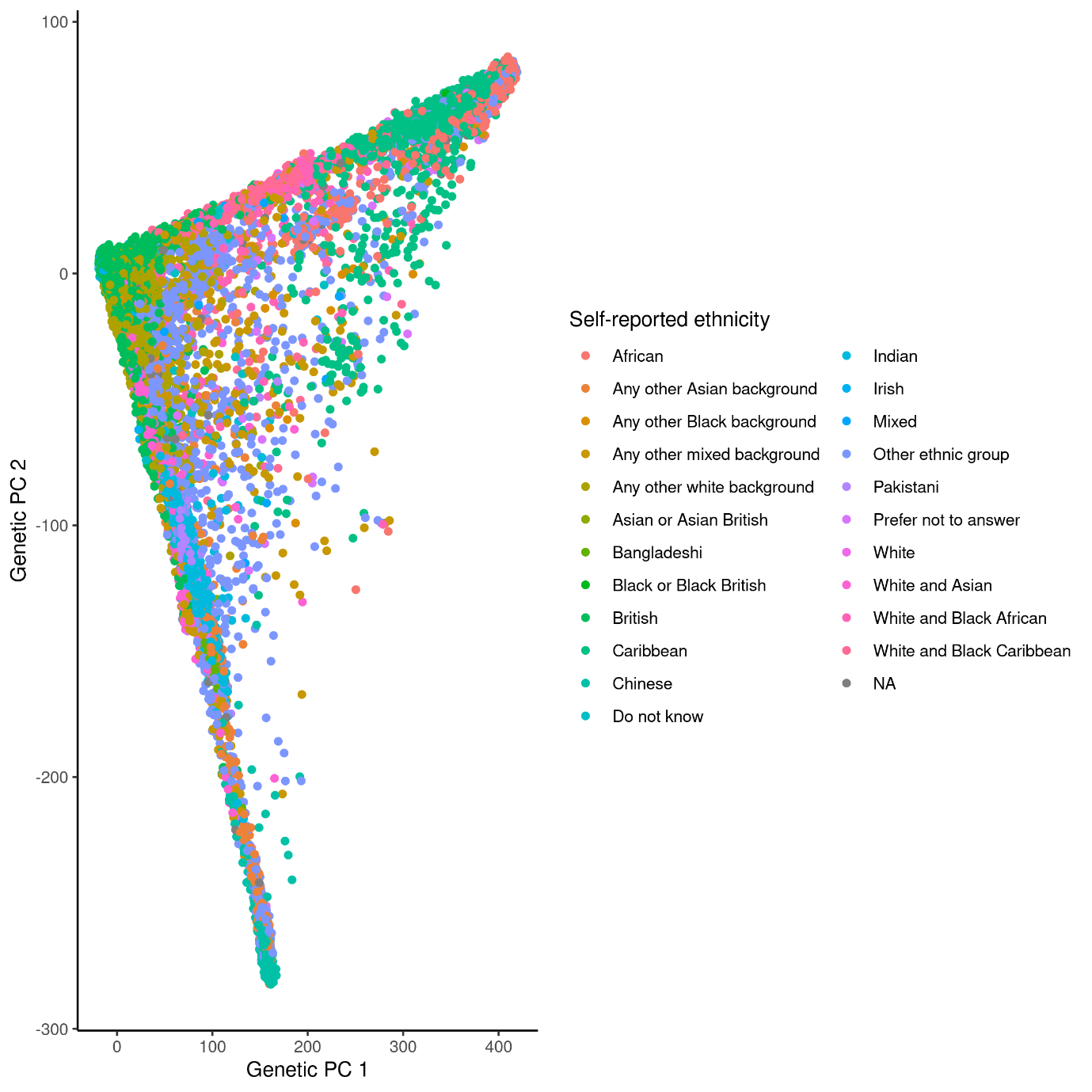

Supplementary figure 4: Genetic principal component plot depicting PCs 1 and 2 after ancestry exclusions. Self-reported ethnicity is shown.

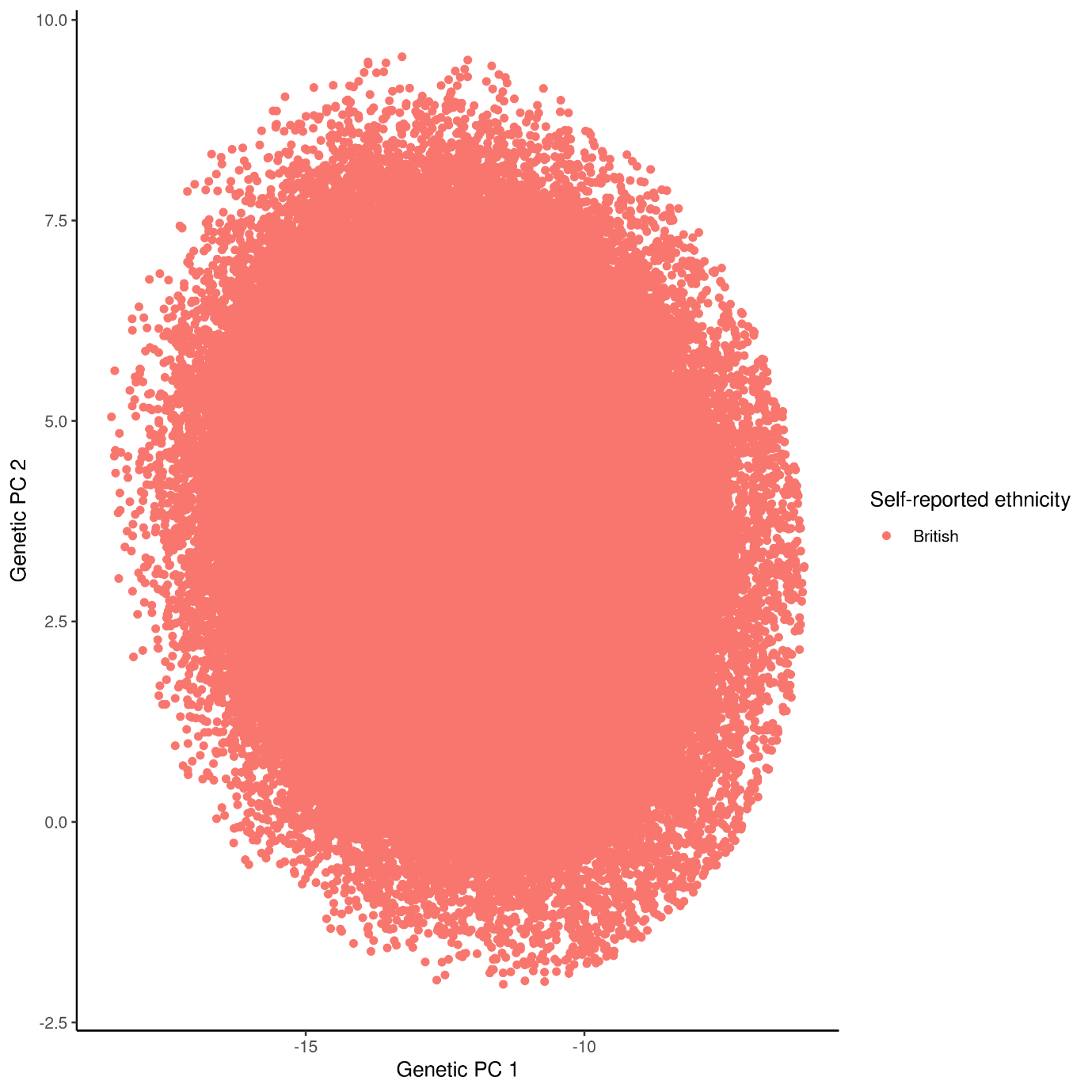

Supplementary figure 5: Nagelkerke’s Pseudo-R2 metric for all 32 MHC PRS without controlling for principal components.

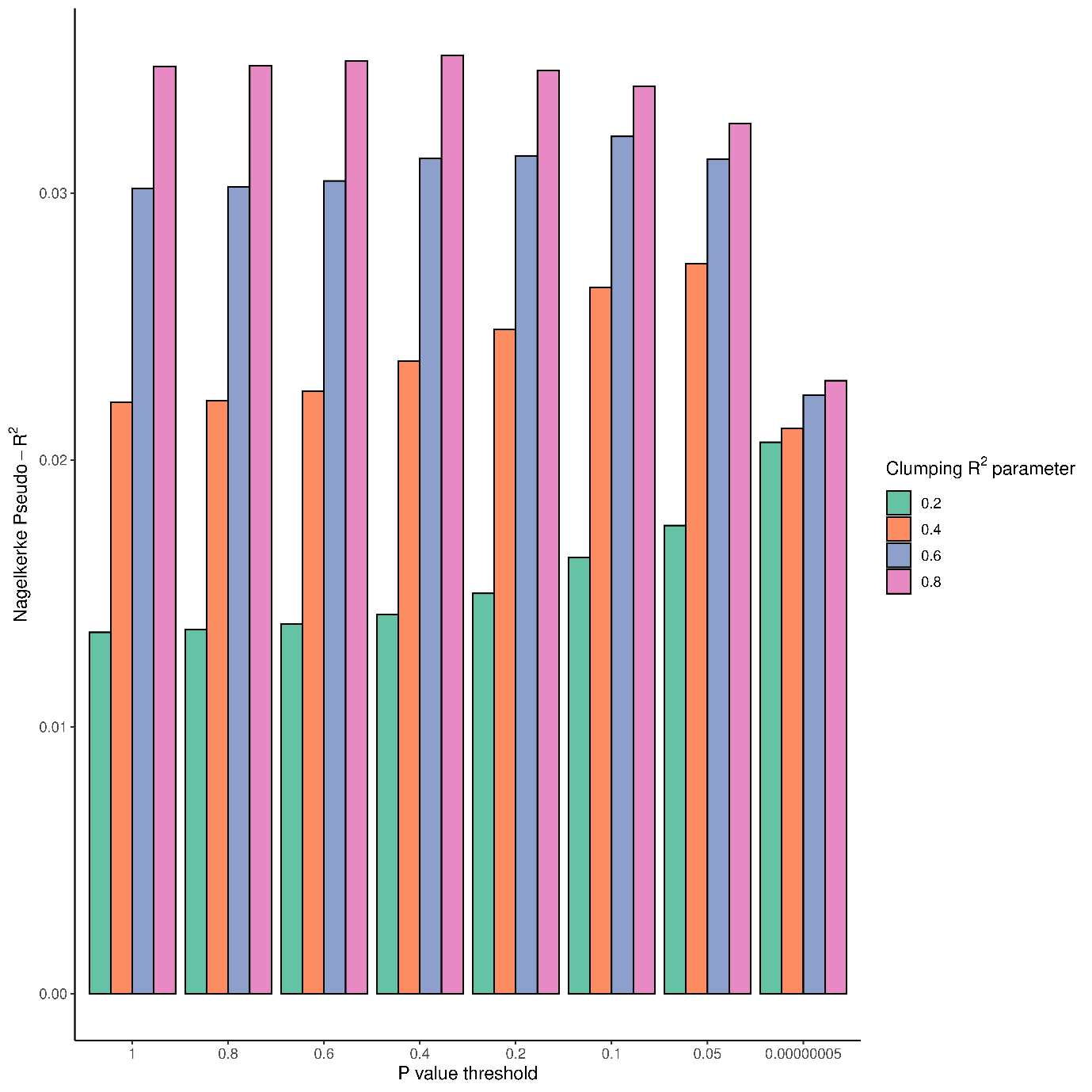

Supplementary figure 6: Nagelkerke’s Pseudo-R2 metric for all 32 MHC PRS controlling for ten principal components.

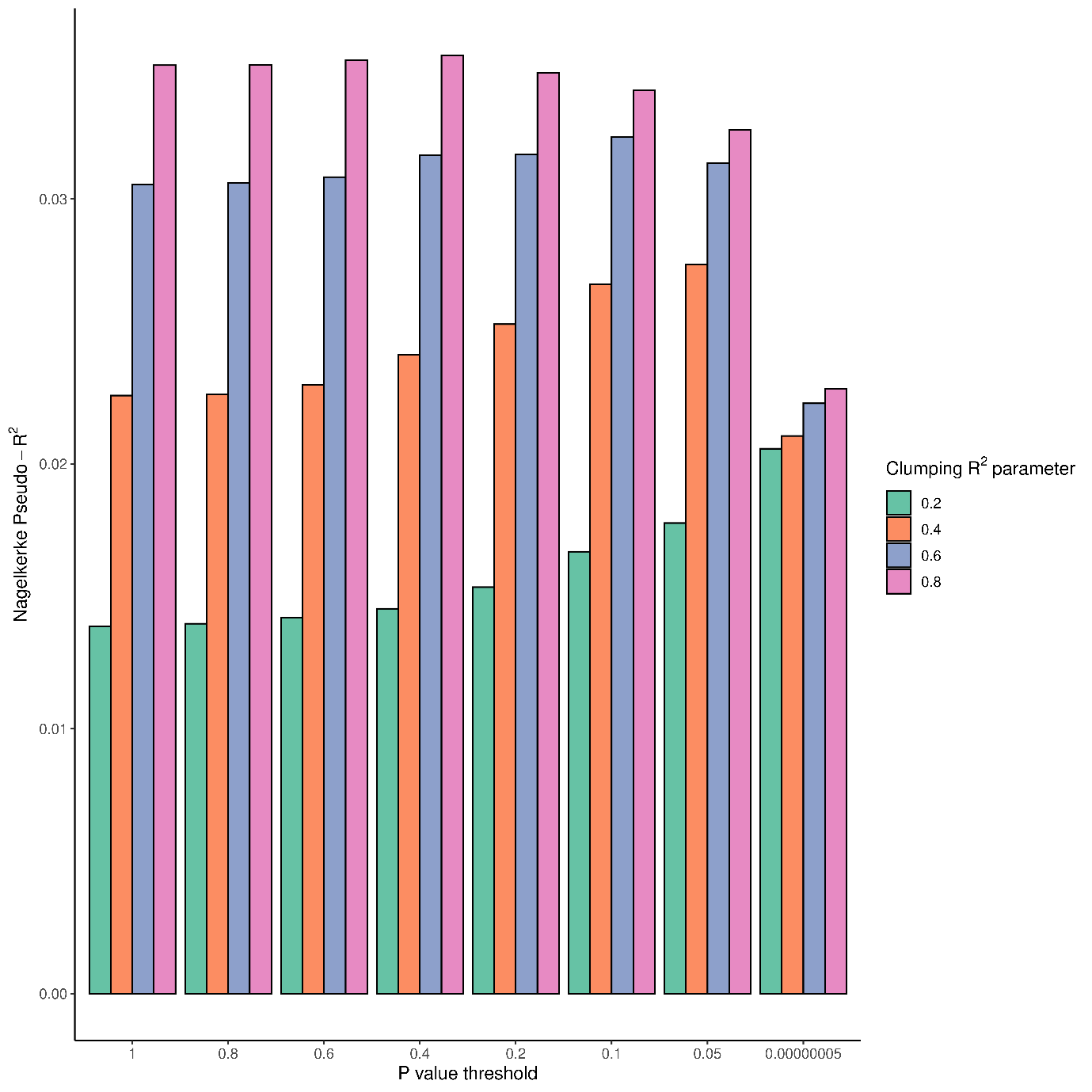

Supplementary figure 7: no association between PRS and claiming of disability benefits among people with MS.

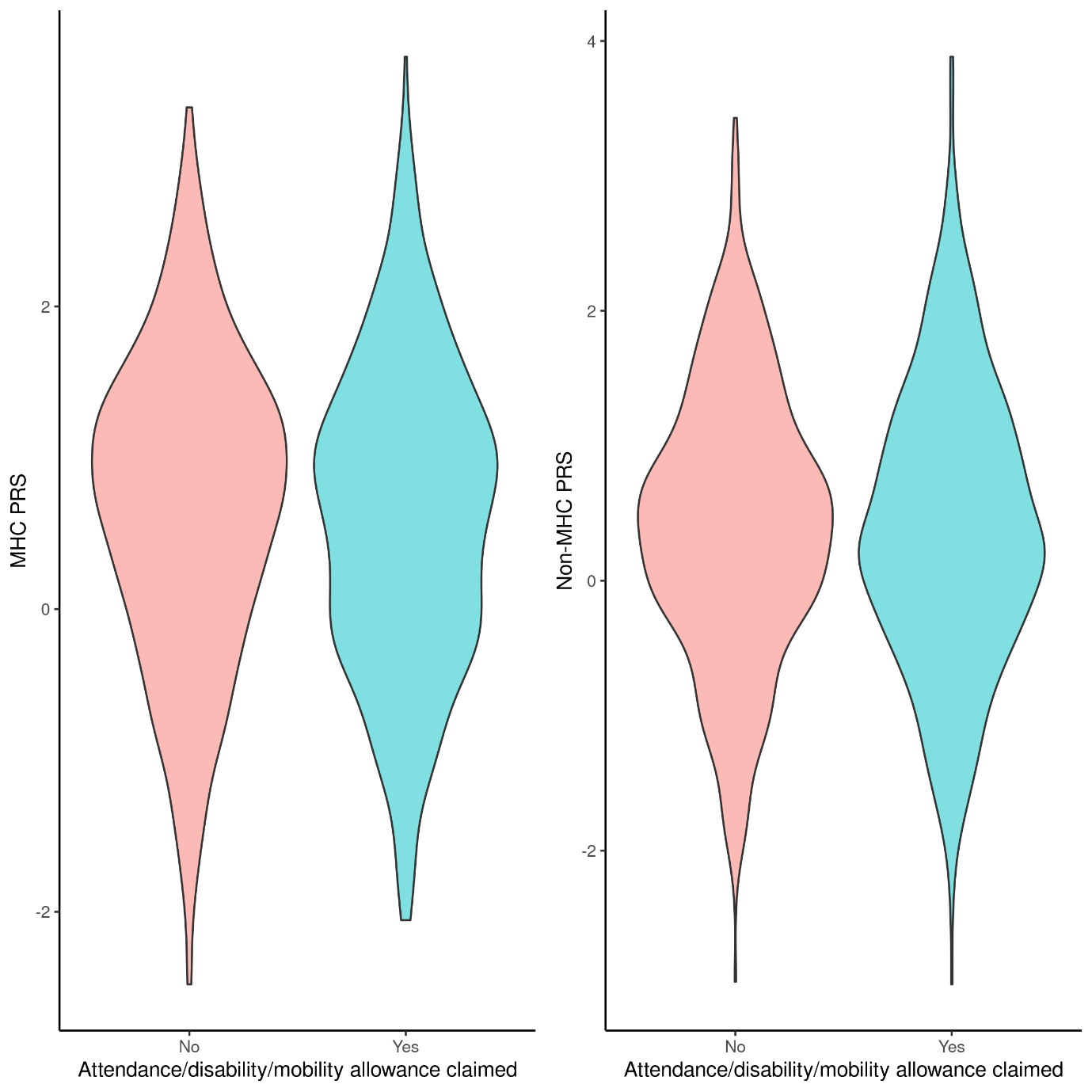

Supplementary figure 8: multiplicative interaction terms and 95% confidence intervals for interaction between environmental exposures and MS genetic risk (MHC and non-MHC scores).

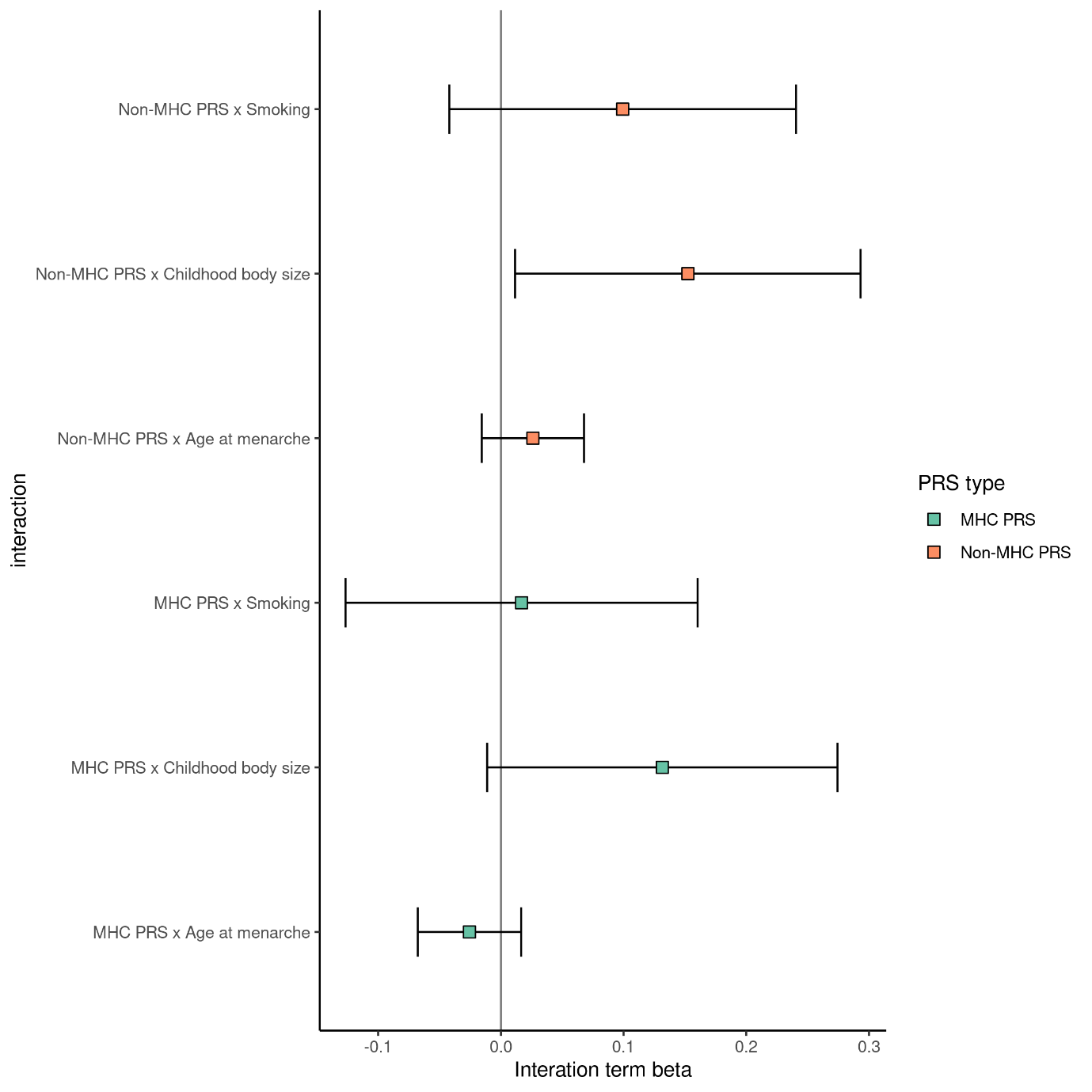
